# Design and Characterisation of Photoactivatable and Lysine Reactive *o*-Nitrobenzyl Alcohol-Based Crosslinkers

**DOI:** 10.1101/2025.05.06.652371

**Authors:** Adam Cahill, Martin Walko, Benjamin Fenton, Sri Ranjani Ganji, Anne Herbert, Sheena E. Radford, Nikil Kapur, Keith Livingstone, Megan H. Wright, Antonio N. Calabrese

**Affiliations:** Astbury Centre for Structural Molecular Biology, School of Molecular and Cellular Biology, Faculty of Biological Sciences, University of Leeds, LS2 9JT, Leeds, United Kingdom; Astbury Centre for Structural Molecular Biology, School of Chemistry, Faculty of Engineering and Physical Sciences, University of Leeds, LS2 9JT, Leeds, United Kingdom; School of Mechanical Engineering, Faculty of Engineering and Physical Sciences, University of Leeds, Leeds LS2 9JT, U.K

## Abstract

Photoreactive groups are invaluable tools in structural proteomics, offering reagent-free activation and temporal control of protein labelling. However, traditional UV-activatable functional groups often produce unstable intermediates and diverse products, making these chemistries difficult to deploy at scale. In this study, we performed a systematic analysis of *ortho*-nitrobenzyl alcohol (*o*NBA) reactivity for integration into novel reagents for chemical crosslinking-mass spectrometry. *o*NBA photochemistry represents a promising alternative to traditional photoactivatable crosslinkers due to its unique specificity towards lysine residues. Here, we synthesised two molecules comprising *o*NBA functional groups with different substituents and assessed their labelling efficiency against a model protein. To ensure high labelling yields while maintaining a short irradiation time, we constructed a high power 365 nm irradiation device which improves the efficiency of *o*NBA photolysis. Our studies identified an amide-substituted probe that labels proteins with high efficiency. We next incorporated this optimised *o*NBA moiety into a homo-bifunctional crosslinker and a hetero-bifunctional crosslinker in combination with an NHS ester, which both resulted in high yields of crosslinked products. Our findings highlight that optimised *o*NBA-based reactive groups are viable UV-activated warheads that can deliver high labelling yields and efficient protein crosslinking, unlocking a wealth of potential structural proteomics applications.

## Introduction

Crosslinking mass spectrometry (XL-MS) is a well-established technique used to understand protein structure, dynamics and interactions. XL-MS relies on using small molecules with two chemical warheads (head groups) that can form covalent bonds with protein residues under physiological conditions^1–3^. Due to the ability of XL-MS to extract information on the non-covalent interactions a protein can form, many crosslinkers have been synthesised utilising different chemistries, to maximise the structural information that can be gathered from XL-MS datasets^4–8^.

Traditional chemical crosslinkers typically target lysine residues due their high intrinsic nucleophilicity and prevalence. The majority of crosslinkers are homo-bifunctional and comprise two *N*-hydroxy succinimide esters (NHS-esters) as lysine-reactive groups, which react with high efficiency under physiological conditions^9^. However, due to the NHS-ester being constitutively active, reactions are always initiated at the point of crosslinker addition. As a result, there is very little temporal control of the crosslinking reaction. This also renders NHS-ester based crosslinkers liable to hydrolysis, forming unproductive ‘mono-links’ (where one of the reactive groups is hydrolysed into an acid that can no longer react with a lysine side-chain). Due to the uncertainty of whether or not the hydrolysis of an NHS-ester has occurred before or after the other NHS-ester had reacted with lysine, mono-links do not deliver the same level of structural information as crosslink^10^.

The ability to temporally control crosslinking using reagent free activation of reactive groups (e.g. via UV irradiation) may enable XL-MS to deliver a better description of the dynamics in protein structures and protein-protein interactions. For example, diazirines and benzophenones have gained traction as light-activatable chemistries incorporated into crosslinker designs ^11,12^. Both head groups are inert under physiological conditions but after UV irradiation form highly reactive species that rapidly insert into proximal covalent bonds ^13^. This not only allows crosslinking reactions to be initiated at chosen time points but also ensures that all the crosslinks formed describe the protein structure(s) present within a time frame determined by the lifetime of the reactive species. However, the generation of unstable intermediates increases the potential number of crosslinks, as the activated species are less selective and may react with many residue sidechains ^14^. This makes analysis of the resultant XL-MS data more computationally expensive, and the precise residue-level localisation of crosslinks can become more difficult. Additionally, as both diazirines and benzophenones can react with water, crosslinking reagents can be ‘quenched’ by solvent and form ‘mono-links’ in analogy with crosslinkers based on NHS-esters.

Given the shortcomings of current crosslinkers, additional chemistries for protein crosslinking are urgently needed to enable the diversity of potential biological applications of XL-MS. Using UV-activatable species that also have high reaction specificity could address these drawbacks, increasing crosslinking yields and simplifying searches to allow easier detection of time-resolved crosslinks. *Ortho*-nitrobenzyl alcohol (*o*NBA) chemistry offers a promising alternative in this regard due to its specific reactivity with lysine residues ^15,16^. Upon UV irradiation, the *o*NBA moiety undergoes dehydration, forming an *o*-nitroso benzaldehyde that is reactive towards lysine sidechains and can undergo a reversible reaction with water (**Figure 1**)^17^. Reports have recently demonstrated the potential of this motif by using *o*NBA crosslinkers to map time-dependent protein-protein interactions using visible light ^18^. However, previous attempts at creating UV-activated *o*NBA based crosslinkers have results in poor yields, unpredictable labelling chemistry and lengthy irradiation times ^19^. To establish *o*NBA as a viable alternative to existing reactive groups, these issues must be addressed. In this work, we report a comparative study between homo- and heterobifunctional crosslinkers containing different *o*NBA and NHS-ester warheads. Our optimised crosslinkers exhibited improved reactivity towards lysine residues in proteins and our workflow exploited a novel UV irradiation platform to deliver high crosslinking yields with < 1 min of irradiation. We further characterised the reactivity of the *o*NBA moieties for biological applications to demonstrate their long-lived active state. Our findings suggest that *o*NBA reactive groups could offer unique advantages over current chemical warheads used in XL-MS and are ideal candidates for time-resolved studies to inform on the structures/rearrangements of the transient molecular assemblies that underpin biological processes.

**Figure 1.**
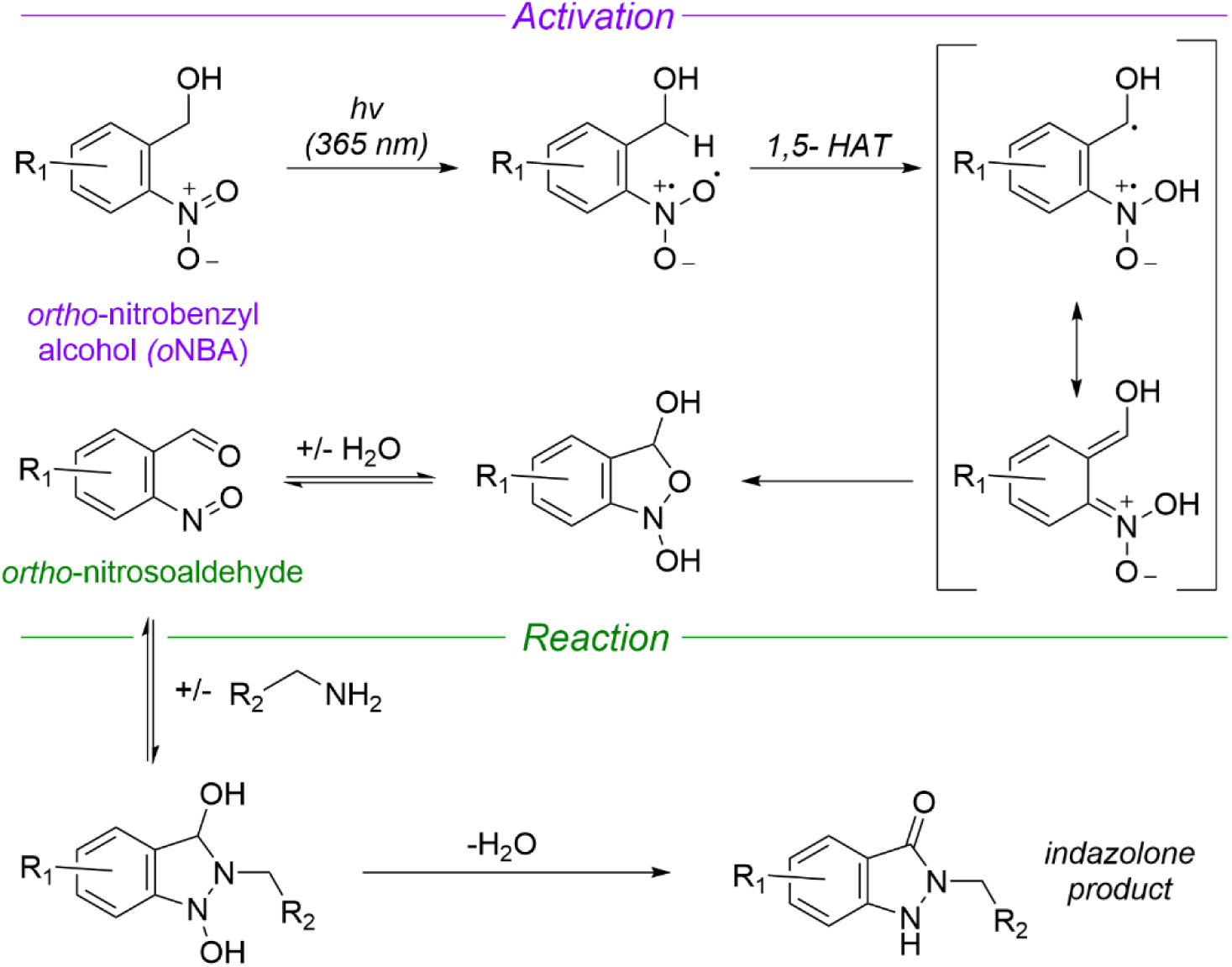
Mechanism of UV-induced activation and reaction of ortho nitrobenzyl alcohols (oNBA) with primary amines. Reproduced from ref.^20^ and ref.^16^. *o*NBA undergoes a light mediated dehydration to form an *ortho*-nitrosoaldehyde. The subsequent reaction terminates with an irreversible condensation to form an indazolone ring (1,5-HAT = 1,5-hydrogen atom transfer).

## Experimental

### Organic synthesis

All reactions were conducted under an atmosphere of nitrogen and protected from light using aluminium foil. All solvents were purchased from Fisher Scientific at HPLC grade and reagents were purchased from Sigma-Aldrich, Fluorochem, Thermofisher and Merck Millipore. Analytical thin layer chromatography (TLC) was conducted using Merck 0.25 mm silica gel pre-coated aluminium plates (Merck silica 2880 gel, 60 F254) and visualised using UV-light. Flash column chromatography was carried out with silica gel 60 (35-75 µm particles). Solvents were removed using a Büchi rotary evaporator and a Vacuubrand PC2001 Vario Diaphragm pump. ^1^H and ^13^C NMR spectra were acquired on a Bruker AV-NEO NMR spectrometer at 500 MHz for ^1^H or 101 MHz for ^13^C. Chemical shifts are expressed as parts per million using solvent as an internal standard (DMSO-*d_6_* 2.50 ppm in ^1^H and 39.52 ppm in ^13^C spectra) and coupling constants are expressed in Hz. The following abbreviations are used: s for singlet, d for doublet, t for triplet, q for quartet, p for pentet, m for multiplet and br for broad. NMR spectra can be found in the **Supplementary Information**.

### Synthesis of 4-(hydroxymethyl)-3-nitro-N-2-propyn-1-ylbenzamide (1a)

4-(Hydroxymethyl)-3-nitrobenzoic acid (100 mg, 0.507 mmol), 1-(3-Dimethylaminopropyl)-3-ethylcarbodiimide hydrochloride (150 mg, 0.783 mmol) and Hydroxybenzotriazole (135 mg, 1 mmol) were dissolved in CH_2_Cl_2_ (20 mL). The reaction was stirred at room temperature for 30 mins before propargylamine (220 µL, 3.4 mmol) and *N,N*-diisopropylethylamine (1 mL, 5.74 mmol) were added. The reaction was left stirring overnight at room temperature. The solvent was then removed *in vacuo* to yield a brown solid. The crude product was purified by silica gel chromatography (1:1 ethyl acetate:hexane) and collected fractions were concentrated to produce a pale yellow solid (77 mg, 0.329 mmol, 66%, R_f_: 0.25 (1:1 ethyl acetate:hexane)).

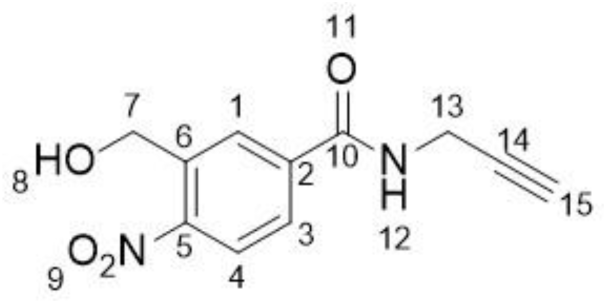

^1^H NMR (500 MHz, DMSO-*d_6_*) δ/ppm 3.17 (t, *J* = 2.5 Hz, 1H, H-C_15_), 4.09 (dd, *J* = 2.5, 5.5 Hz, 2H, H-C_7_), 4.88 (d, *J* = 5.5 Hz, 2H, H-C*13*), 5.67 (t, *J* = 5.5 Hz, 1H, OH), 7.95 (d, *J* = 8.1 Hz, 1H, H-C_4_), 8.23 (dd, *J* = 1.8, 8.1 Hz, 1H, H-C_3_), 8.53 (d, *J* = 1.8 Hz, 1H, H-C_1_), 9.27 (t, *J* = 5.4 Hz, 1H, NH). ^13^C NMR (101 MHz, DMSO-*d_6_*) δ/ppm: 28.7 (C_13_), 59.9 (C_7_), 73.2 (C_14/15_), 80.9 (C_14/15_), 123.2 (C_Ar_), 128.6 (C_Ar_), 132.2 (C_Ar_),133.2 (C_Ar_), 141.7 (C_Ar_), 146.6 (C_Ar_), 163.8 (C_10_).

### Synthesis of (2-Nitro-5-(prop-2-yn-1-yloxy)phenyl)methanol (1b)

To 5-hydroxy-2-nitrobenzyl alcohol (169 mg, 0.998 mmol) and anhydrous potassium carbonate (2.76 mg, 0.019 mmol) was added in 1.5 mL of anhydrous DMF and the resulting mixture was stirred for 2 hours at 60°C. Then propargyl bromide (0.15 mL, 2 mmol) was slowly added into the reaction and the orange mixture was stirred for 24 hours at room temperature. After the reaction was complete the solvent was removed under reduced pressure and the residue was resuspended in ethyl acetate (10 mL). The suspension was then washed once with water (10 mL) and three times with brine (10 mL). After this the organic layer was dried over MgSO_4_ and filtered. The solvent was then removed under reduced pressure and the residue purified via flash column chromatography on silica gel (1:1 ethyl acetate:hexane). The sample was concentrated until dryness to deliver a yellow solid (123 mg, 0.594 mmol, 59%, R_f_ 0.34 (1:1 ethyl acetate:hexane)).

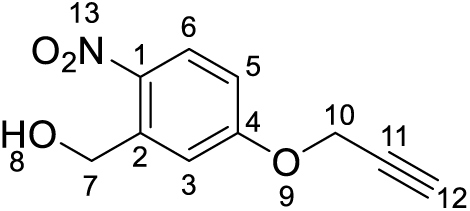

^1^H NMR (500 MHz, DMSO-*d_6_*) δ/ppm: 3.67 (t, *J* = 2.4 Hz, 1H, H-C_12_), 4.85 (d, *J* = 5.5 Hz, 2H, H-C_7_), 4.97 (d, *J* = 2.4, 2H, H-C_10_), 5.59 (t, *J* = 5.5 Hz, 1H, H-C_8_), 7.09 (dd, *J* = 2.9, 9.1 Hz, 1H, H-C_5_), 7.41 (d, *J* = 2.9 Hz, 1H, H-C_3_), 8.15 (d, *J* = 9.1 Hz, 1H, H-C_6_). ^13^C NMR (101 MHz, DMSO-*d*_6_) δ/ppm: 56.14 (C_7_), 60.1(C_10_), 78.4 (C_12_), 79.1 (C_11_), 113.1 (C_Ar_), 113.8 (C_Ar_), 127.4 (C_Ar_), 139.9 (C_Ar_), 142.2 (C_Ar_), 161.5 (C_Ar_).

### Synthesis of 2,5-dioxo-1-pyrrolidinyl 4-(hydroxymethyl)-3-nitrobenzoate (2)

4-(Hydroxymethyl)-3-nitrobenzoic acid (100 mg, 0.507 mmol), *N*-hydroxysuccinimide (120 mg, 1.043 mmol) and 1-(3-dimethylaminopropyl)-3-ethylcarbodiimide hydrochloride (125 mg, 0.805 mmol) were added to a round bottom flask. DMF (5 mL) was added, and the resulting orange solution was stirred overnight at room temperature. The reaction mixture, deionised water (20 mL) and ethyl acetate (20 mL) were added to a separating funnel and phases were separated. The aqueous layer was removed, and the organic layer was washed three times with water (20 mL) and then dried over MgSO_4_, filtered and concentrated to yield a yellow solid. The crude product was purified by silica gel chromatography (2:1 ethyl acetate:hexane) to yield the title compound as a white-yellow solid (80 mg, 0.272 mmol, 53%, R_f_: 0.28 (2:1 ethyl acetate:hexane)).

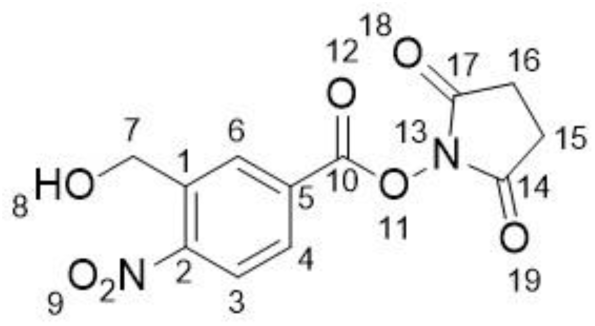

^1^H NMR (500 MHz, DMSO-*d_6_*) δ/ppm: 2.91 (s, 4H, H-C_15_H-C_16_), 4.96 (d, *J* = 5.5 Hz, 2H, H-C_7_), 5.84 (t, *J* = 5.5 Hz, 1H, H-C_8_), 8.15 (d, *J* = 8.2 Hz, 1H, H-C_3_), 8.46 (dd, *J* = 1.8, 8.2 Hz, 1H, H-C_4_), 8.63 (d, *J* = 1.8 Hz, 1H, H-C_6_). ^13^C NMR (101 MHz, DMSO-*d_6_*) δ/ppm: 25.6 (C_15_C_16_), 60.0 (C_7_), 123.8 (C_Ar_), 125.9 (C_Ar_), 129.7 (C_Ar_), 134.5 (C_Ar_), 146.2 (C_Ar_), 146.9 (C_Ar_), 160.4 (C_10_), 170.1 (C_14_C_17_).

### Synthesis of *N*,*N*′-1,2-ethanediylbis(3-hydroxymethyl-4-nitrobenzamide (3)

4-(hydroxymethyl)-3-nitrobenzoic acid (290 mg, 1.47 mmol), 1-(3-dimethylaminopropyl)-3-ethylcarbodiimide hydrochloride (325 mg, 1.69 mmol), and *N*-Hydroxybenzotriazole (230 mg, 1.69 mmol) was added to a flask and flushed with N_2_. The mixture was then dissolved in acetonitrile (20 mL) and left to stir for 30 mins at room temperature. Ethylenediamine was then added (270 µL, 0.43 mmol) as well as DIPEA (300 µL, 1.69 mmol) and the reaction was left to stir overnight at room temperature. Once the reaction was complete, the mixture was gravity filtered and the solution was dried *in vacuo*. The crude product was dissolved in 10 mL of 1:1 mixture of acetonitrile and water and purified using preparative HPLC on an Agilent 1260 infinity system equipped with UV detector and fraction collector on Kinetex EVO 5 um C18 100A 21.2 x 250 mm reverse phase column. The solution (4.5 mL) was injected, and a 25 min gradient of 20 – 50 % acetonitrile in water with 0.1% formic acid was performed at 15 mL/min. The fractions containing the product were combined and freeze dried to afford the title compound as a white solid (10 mg, 24 µmol, 6%).

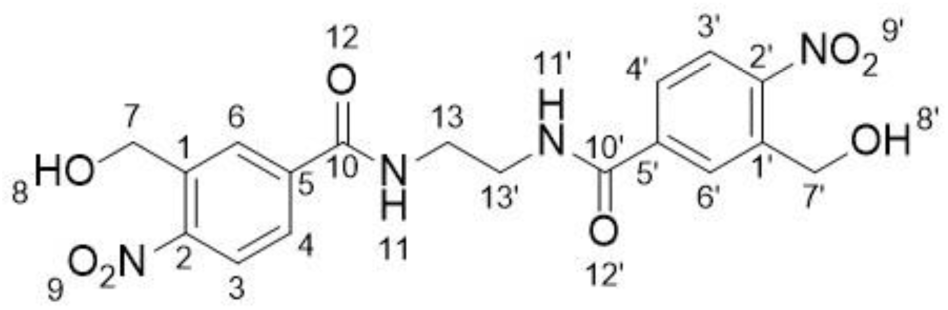

^1^H NMR (500 MHz, DMSO-*d_6_*) δ/ppm 3.43 – 3.56 (m, 4H, H-C_13_), 4.87 (d, *J* = 5.5 Hz, 4H, H-C_7_), 5.65 (t, *J* = 5.5 Hz, 2H, H-C_8_), 7.93 (d, *J* = 8.1 Hz, 2H, H-C_3_), 8.21 (dd, *J* = 1.8, 8.1 Hz, 2H, H-C_4_), 8.51 (d, *J* = 1.8 Hz, 2H, H-C_6_), 8.93 (s, 2H, 2xNH). ^13^C NMR (101 MHz, DMSO-*d*_6_) δ/ppm: 60.4 (C_7_), 123.6 (C_Ar_), 128.9 (C_Ar_), 132.6 (C_Ar_), 134.5 (C_Ar_), 141.8 (C_Ar_), 147.0 (C_Ar_), 164.8 (C_10_).

### SurA expression and purification

This protocol was adapted from previous work ^21^. A pET28b plasmid containing the mature SurA sequence (without the signal peptide sequence) preceded by an *N*-terminal 6x His-tag and a TEV-cleavage site was transformed into BL21(DE3) cells (Stratagene). Cells were grown in LB medium supplemented with 30 µg/mL kanamycin at 37 °C with shaking (200 rpm) until an OD_600_ of ∼0.6 was reached. The temperature was subsequently lowered to 20 °C, and expression induced with 0.4 mM IPTG. After ∼18 h, cells were harvested by centrifugation (4000 xg, 15 min), resuspended in 25 mM Tris-HCl, pH 7.2, 150 mM NaCl, 20 mM imidazole, containing EDTA-free protease inhibitor tablets (Roche), and lysed using a cell disrupter (Constant Cell Disruption Systems). The cell debris was removed by centrifugation (20 min, 4 °C, 39,000 × g), and the lysate was applied to a 5 mL HisTrap column (GE Healthcare). The column was washed with 25 mM Tris-HCl, pH 7.2, 150 mM NaCl and 20 mM imidazole, and SurA was eluted with 25 mM Tris-HCl, 150 mM NaCl, 500 mM imidazole, pH 7.2. The eluate was dialysed against 25 mM Tris-HCl, 150 mM NaCl, pH 8.0 overnight, and the following day TEV protease (ca. 0.5 mg) and 0.1% (v/v) β-mercaptoethanol were added. The cleavage reaction was left to proceed overnight at 4 °C on a tube roller. The cleavage reaction was again applied to a 5 mL HisTrap column (GE Healthcare) to remove the cleaved His-tag and His-tagged TEV protease. The unbound, cleaved SurA product was dialysed against 25 mM Tris-HCl, 150 mM NaCl, pH 8.0, before being concentrated to ∼200 µM with Vivaspin 20 concentrators (Sartorius; 5-kDa MWCO), aliquoted, snap-frozen in liquid nitrogen and stored at −80 °C.

### OmpX expression and purification

OmpX was expressed and purified using a method adapted from ref. ^21^. Briefly, E. coli BL21[DE3] cells (Stratagene) were transformed with a pET11a plasmid containing the gene sequence of OmpX. Overnight cultures were subcultured and grown in LB medium (500 mL) supplemented with carbenicillin (100 μg/mL), at 37 °C with shaking (200 rpm). Protein expression was induced with IPTG (1 mM) once an OD_600_ of 0.6 was reached. After 4 h the cells were harvested by centrifugation (5000 × g, 15 min, 4 °C). The cell pellet was resuspended in 50 mM Tris-HCl pH 8.0, 5 mM EDTA, 1 mM phenylmethylsulfonyl fluoride, 2 mM benzamidine, and the cells were subsequently lysed using a cell disrupter (Constant Cell Disruption Systems). The lysate was centrifuged (25,000 × g, 30 min, 4 °C) and the insoluble material was resuspended in 50 mM Tris-HCl pH 8.0, 2% (v/v) Triton X-100, before being incubated for 1 h at room temperature, with gentle agitation. The insoluble material was pelleted (25,000 × g, 30 min, 4 °C) and the inclusion bodies were washed twice by resuspending in 50 mM Tris-HCl pH 8.0 followed by incubation for 1 h at room temperature with gentle agitation, and then collected by centrifugation (25,000 × g, 30 min, 4 °C). The inclusion bodies were solubilised in 25 mM Tris-HCl, 6 M Gdn-HCl, pH 8.0 and centrifuged (20,000 × g, 20 min, 4 °C). The supernatant was filtered (0.2 µM syringe filter, Sartorius) and the protein was purified using a Superdex 75 HiLoad 26/60 gel filtration column (GE Healthcare) equilibrated with 25 mM Tris-HCl, 6 M Guanidine-HCl, pH 8.0. Peak fractions were concentrated to ∼100 μM using Vivaspin 20 (5 kDa MWCO) concentrators (Sartorius), and the protein solution was snap-frozen in liquid nitrogen and stored at −80 °C.

### UV-Vis absorbance

Compounds ***1a*** and ***1b*** were dissolved in methanol to 25 µg/mL. The absorbance of the molecules was measured across the range 220 nm – 400 nm using a quartz cuvette (1 cm path length) using a UV-1900i spectrophotometer (Shimadzu).

### Construction of UV lamp

The UV lamp was built using a 365 nm fReactor PhotoFlow module (Asynt), which houses the LED, an aspheric condenser lens and cooling fan. A lens tube (Thorlabs - SM1L20) fitted with an aspheric condenser lens (ThorLabs – ACL2520U-A) was mounted onto the module, such that the combined pair of lenses focusses the beam to a spot. The sample tube sits within the lens tube, positioned so that the fluid is located at this spot (see **Figure 2C**). To prevent exposure of the user to UV, the top of the lens tube was fitted with a pair of cage plates (ThorLabs – CP33T/M and CP31/M); the bored cage plate was fitted permanently to the lens tube, and the solid cage plate could be lifted off (also deactivating the lamp). Magnets were used to hold this in place.

**Figure 2.**
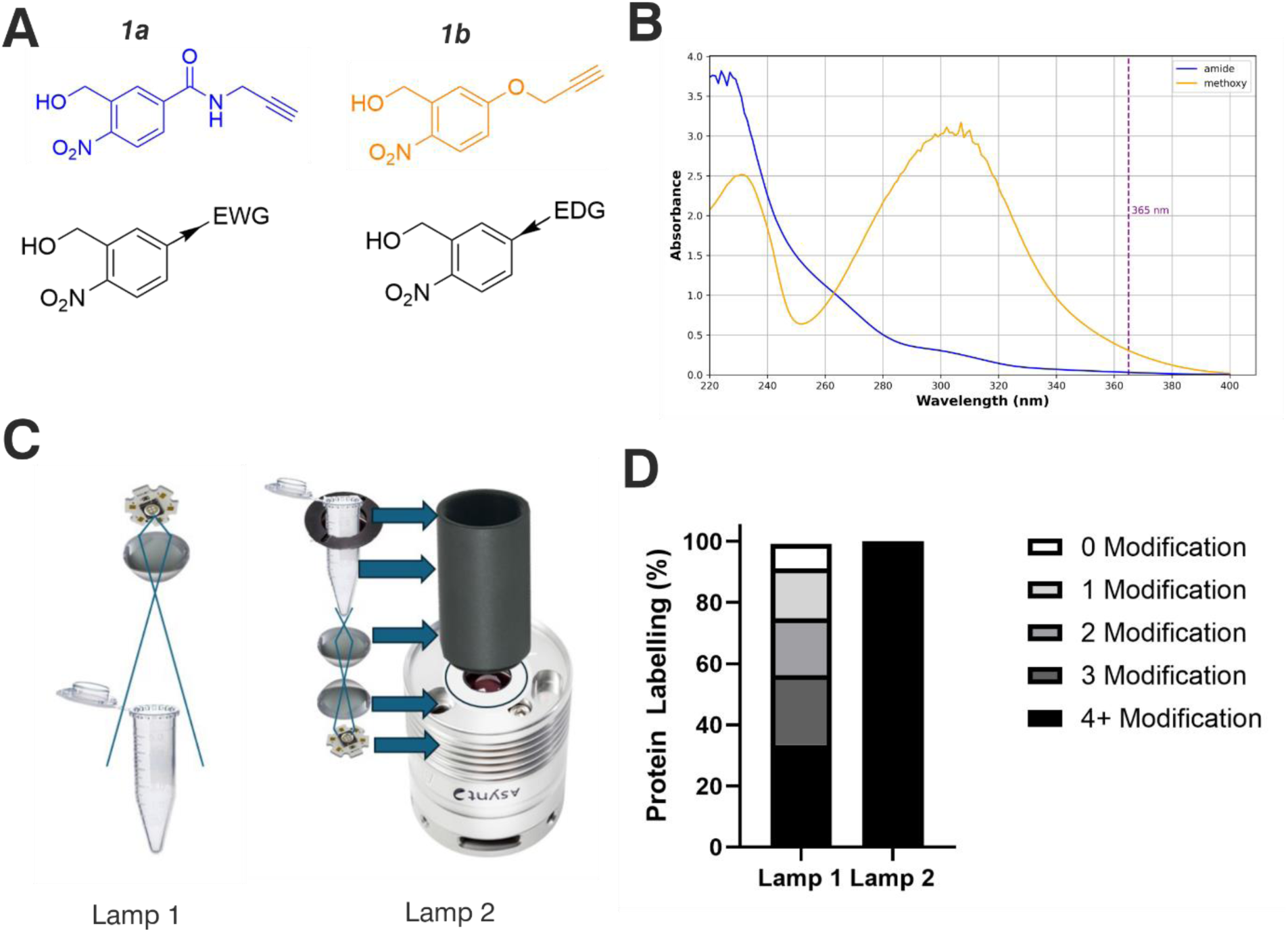
Optimising the physicochemical properties of oNBA warheads for protein labelling and development of an improved high-powered irradiation device to increase labelling efficiency. **(A)** The two synthesised reactive oNBA reactive head groups containing a para-electron withdrawing group (EWG, amide) (**1a**) or electron donating group (EDG, methoxy) (**1b**). **(B)** UV/Vis spectra showing the absorbance of probe **1a** and **1b,** highlighting the lack of absorbance of both oNBA groups at 365 nm. **(C)** Schematics highlighting the structural differences between the two in-house built lamps. For details of lamp 1 see ref. ^28^, and for details of lamp 2 see Methods. **(D)** Protein labelling reactions of SurA using probe **1a**. SurA was incubated with **1a** at a 50-molar excess in PBS with a 30 second UV irradiation time using the two different lamps. The samples were left after irradiation for an hour and then analysed via intact MS. The relative abundance of the differentially labelled species was quantified.

### Protein labelling reactions

SurA was buffer exchanged into phosphate buffered saline (PBS) (137 mM NaCl, 2.7 mM KCl, 10 mM Na_2_HPO_4_, and 1.8 mM KH_2_PO_4_, pH 7.4) using Zeba Spin Columns (Thermo Fisher Scientific) according to the manufacturer’s instructions. The concentration of the protein solution was determined spectrophotometrically (ԑ = 29450 cm^-1^) and the solution was diluted to 20 µM. Both synthesized head groups (***1a*** and ***1b***) were dissolved in methanol at a concentration of 6.25 mM and this was added to the SurA solutions to afford a final concentration of 0.2 - 1 mM. After the addition of the head group, samples (20 µL) containing *o*NBA reactive moieties were irradiated with 365 nm light for 30 seconds.

The samples were then left at room temperature for 1 hour and then then mixed with 0.2 % formic acid (10 µL).

### Intact mass spectrometry

Protein desalting and intact mass analysis was performed by LC-MS using a Vanquish UPLC (Thermo Scientific) interfaced to a Orbitrap Exploris 240 mass spectrometer (Thermo Scientific). Samples were diluted to 10 µm using 0.1% TFA. 3 µL of the sample was injected onto an MAbPac RP column (4 µm, 2.1 x 100 mm, Thermo Scientific). Buffer A was 0.1% (v/v) formic acid in water, and buffer B was 0.1% (v/v) formic acid in acetonitrile. The system flowrate was kept constant at 0.25 ml/min. The bound protein was eluted by a gradient of 20-45% solvent B in A over 1 min and the solvent conditions were then held at 45% solvent B in A for 1.5 min. The column was subsequently washed with a gradient of 45%-80% solvent B over 1.5 min, held at 80% solvent B for 2 min before re-equilibration at 20% solvent B in A ready for the next injection. Data were processed using UniDec ^22^.

### Crosslinking BSA using the hetero-bifunctional crosslinker 3

Bovine Serum Albumin (BSA) (Sigma) was dissolved in PBS and samples were diluted to 20 µM. The synthesized crosslinker **3** (2,5-dioxo-1-pyrrolidinyl 4-(hydroxymethyl)-3-nitrobenzoate) was dissolved in DMSO to a concentration of 25 mM. The crosslinker was added to the protein (final concentration of 2 mM) and the solution was left to incubate at room temperature for 30 mins. The sample was then irradiated with UV light for 30 seconds before another 30 min incubation. The excess crosslinker was then quenched with 2 µL of lysine (1M lysine-HCl, pH 7.4).

### Crosslinking BSA using the homo-bifunctional crosslinker 4

BSA was dissolved in PBS (137 mM NaCl, 2.7 mM KCl, 10 mM Na_2_PO_4_, 1.8 mM K_2_PO_4_, pH 7.4) and samples were diluted to 20 µM. The synthesized crosslinker **4** (N,N′-1,2-ethanediylbis(3-hydroxymethyl-4-nitrobenzamide), was dissolved in DMSO to a concentration of 25 mM. The crosslinker was added to 20 µL of protein (final concentration of 2 mM) and the samples were irradiated with UV light for 30 seconds before a 30 min incubation. The excess crosslinker was then quenched with 2 µL of lysine (1M lysine-HCl, pH 7.4).

### Sodium dodecyl sulphate – polyacrylamide gel electrophoresis

Samples were diluted 3:1 with 4x loading buffer (100 mM Tris-HCl pH 6.8, 200 mM DTT, 4% (w/v) SDS, 0.2% (w/v) bromophenol blue, 20% (w/v) glycerol) and then boiled for ten minutes. Boiled samples were spun down in a bench-top centrifuge for one minute and 10 µL of sample was loaded into wells of a Mini-PROTEAN 12% pre-cast gels (Bio-Rad). Precision plus protein dual colour protein ladder (BioRad, CA, USA) was loaded in the first gel lane as a standard for molecular weight determination (5 µL). Gels were run at a constant current of 60 mA until the dye front reached the bottom of the gel. Gels were removed from casting plates and were stained in InstantBlue Coomassie protein Stain overnight and imaged on an Alliance Q9 imager (Uvitec).

### Digestion of proteins

Cross-linked proteins were digested using S-Trap micro columns (Protifi), according to the manufacturer’s instructions using MS-grade solvents. First, 25 µL of 10 % (w/v) SDS solution was added to 25 µL of crosslinked protein (concentration of 20 µM). DTT solution (5 µL, 220 mM) was added and the sample was heated to 95°C for 15 mins and then allowed to cool to room temperature. Iodoacetamide (5.5 µL, 440 mM) was added and the sample was kept in the dark for 30 mins at room temperature. The solution was acidified by adding phosphoric acid to a final concentration of 1.2 % (v/v). Samples were then diluted with S-Trap binding buffer 7:1 (100 mM triethylammonium bicarbonate (TEAB) pH 7.1 in methanol). Trypsin (Promega) was added from stocks at a concentration of 0.02 µg/ µL to a ratio of 1:20 (enzyme:protein). The protein solution containing the appropriate enzyme was loaded onto the S-Trap column and centrifuged at 4000 x g for 30 s, to enable protein capture within the submicron pores of the S-Trap. The column was washed by adding 130 µL of binding buffer before being centrifuged at 4000 x g for 30 s. 30 µL of trypsin (0.02 µg/ µL in TEAB) was added to the top of the S-trap column. The S-Trap was loosely capped, placed in a 1.5 mL microcentrifuge tube and heated at 47 °C for 90 mins. 40 µL of 50 mM TEAB was added and the column was centrifuged at 4000 x g for 1 min. Digested peptides were eluted by sequential washing with of 0.2 % (v/v) formic acid (FA) and then 50 % (v/v) acetonitrile (ACN), with centrifugation (4000 x g for 30 s) after each addition. The eluates from these steps were combined and dried using a Concentrator Plus vacuum centrifuge (Eppendorf).

### Crosslinking-mass spectrometry

Dried samples were resuspended in 0.1 % (v/v) trifluoracetic acid to a concentration of *ca.* 1 µM. Samples were then analysed on Vanquish Neo LC System (Thermo Scientific) coupled to an Orbitrap Eclipse Tribrid mass spectrometer (Thermo Scientific). The instrument was fitted with an EASY-spray reversed phase LC system with column specifications as follows: particle size: 2 µm, diameter: 75 µm, length: 500 mm, running at 250 nL/min flow and kept at 45°C. Analytes were eluted by a 65 min, 0-50% acetonitrile gradient. The mass spectrometric settings for MS1 scans were: resolution of 120 000, automatic gain control (AGC) target of 3 × 10^6^, maximum injection time of 50 ms, scanning from 380– 1450 m/z in profile mode. Cycle time was adjusted to 3 seconds and charge states z = 3–8 were isolated using a 1.4 m/z window and fragmented by HCD using optimized stepped normalized collision energies, 30 ±6. Fragment ion scans were acquired at a resolution of 60 000, AGC target of 1 × 10^5^, maximum injection time of 120 ms, scanning from 200–2000 m/z, underfill ratio set to 1%. Dynamic exclusion was enabled for 30 s (including isotopes). Thermo RAW files were converted to mgf files using Proteome Discoverer (Thermo Scientific). The mgf files were searched using MeroX to identify crosslinked peptides ^23^. The following settings were applied: proteolytic cleavages C-terminally to lysine and arginine; up to 3 missed cleavages; peptide length 5 to 30 amino acids; modifications: alkylation of cysteine by iodoacetamide, oxidation of methionine; crosslinker specificity lysine; precursor precision 10 ppm; fragment precision 20 ppm; signal to noise >2; FDR cut off 1%. Mapping of crosslinks onto structures and distance measurements (Cα-Cα Euclidean distances) was performed using the PyXlinkViewer plugin for PyMol ^24^.

### Mapping of covalent modifications by mass spectrometry

Dried samples were resuspended in 0.1% (v/v) trifluoracetic acid at a concentration of *ca* 1 µM. Samples were then analyzed on Vanquish Neo LC System (Thermo Scientific) coupled to Orbitrap Eclipse Tribrid mass spectrometer (Thermo Scientific). The instrument was fitted with an EASY-spray reversed phase LC system with column specifications as follows: particle size: 2 µm, diameter: 75 µm, length: 500 mm, running at 250 nL/min flow and kept at 45°C. Analytes were eluted by 65 min, 0-50% acetonitrile gradient. The mass spectrometric settings for MS1 scans were: resolution of 120 00, AGC target of 3 x 10^6^, maximum injections time of 50 ms, scanning from 350-2000 m/z in profile mode. Cycle time was adjusted to 2.5 seconds and charge states z = 2-7 were isolated using a 1.2 m/z window and fragmented by HCD using optimized stepped normalized collision energies, 30 ±6. Fragment ions scans were acquired at a resolution of 30 000, AGC target of 5 x 10^4^, maximum injection time of 54 ms, scanning from 200-2000 m/z underfill ratio set to 1%. Dynamic exclusion was enabled for 60 s (including isotopes). The data were analyzed using PEAKS Studio 11.5 (Bioinformatics Solutions). The reference BSA sequence was retrieved from UniProt (P02769). The enzyme was set to trypsin with specific cleavages and a maximum of two missed cleavages. Fixed modification was set to carbamidomethyl (C: +57.02). The variable modifications were: deamidation (NQ: +0.98), oxidation (M: +15.99), indazolone *o*NBA amide (K: +198.04), secondary amide-*o*NBA amide (K: +200.06), indazolone *o*NBA methoxy (K: +171.03) or secondary amide-*o*NBA methoxy (K: +173.05). The allowed peptide lengths were between 6 – 45. The FDR was set at 1%.

## Results and Discussion

### Design of reactive groups and light source

Studying the fundamental chemical reactivity of crosslinkers is complicated by factors such as the availability of proximal lysine residues, steric restrictions of reactive groups and the possibility of multiple reaction products. Therefore, to characterise and optimise *o*NBA reactivity with proteins effectively, we first synthesised two different *o*NBA reactive head groups that have either amide (electron withdrawing, ***1a***) or methoxy (electron donating, ***1b***) substituents to alter the physicochemical and photophysical properties of the *o*NBA moiety (**Figure 2A**). The *o*NBA groups were coupled to an alkyne due to its inert behaviour in biological applications and potential future use as a handle in chemical biology. We gave careful consideration to the wavelength of light to be used for irradiation due to the potential biological damage that can occur at wavelengths < 300 nm. While *o*NBA moieties have been applied to chemical crosslinking using wavelengths of > 400 nm ^18^, we elected to employ 365nm light that is also commonly used with conventional photoactivatable reagents like diazirines and benzophenones^25,26^, as this would enable our reagents to be widely deployed in the XL-MS community. The increase of π-electron density imparted by addition of the methoxy substituent increased the λ_max_ of probe **1b** to ∼300 nm, with its absorbance at 365 nm increased 10-fold relative to **1a** (**Figure 2B**).

Given the low absorbance of **1a** relative to **1b** at 365 nm, we sought to develop a high-power lamp to increase the rate of head group photolysis to minimise differences in irradiation time^27^. In previous work, we developed a lamp with a 3.8 W UV-LED positioned above the sample ^28^ (**Figure 2C, left**). Here, we developed a new apparatus that utilised a LED with an additional focal lens, positioned below the sample, directing focused light upwards^28^. This design reduced both the path length to the sample and the light cone angle, minimising the loss of light to the surrounding environment and maximising irradiation of the sample. Initial test reactions using the globular protein SurA and probe **1a** showed that these alterations greatly improved the labelling efficiency achievable using this new device, as intact-MS analysis revealed complete modification of SurA after just 30 seconds of irradiation (50 molar excess of **1a**, **Figure 2D**).

### Characterisation of *o*NBA labelling

With the ability to achieve high-yielding labelling in short irradiation times, we further characterised both *o*NBA analogues. We incubated a model globular protein (SurA, 45.1 kDa, 21 lysine residues) with either of the *o*NBA analogues at increasing concentrations (ranging from 1:10 to 1:50 molar ratio of protein:reagent), or an NHS-ester reactive group for comparison. After activation (30 seconds) and an incubation time of 1 hour, we analysed the samples using intact MS to identify and quantify the species present to determine the degree of labelling. Head group **1b** exhibited significantly lower labelling efficiency compared to **1a** despite its higher absorbance at 365 nm (**Figure 3A** and **3B**). The yields of labelling by **1a** were similar to those of the NHS-ester, with both producing highly labelled species (>4 modifications) at all concentrations tested (**Figure 3B** and **3C**). To investigate the kinetics of protein labelling by photoproducts of *o*NBA irradiation, SurA was incubated with **1a**, the samples were irradiated with 365 nm light for 30 seconds, and the labelling reaction was terminated at different time points after irradiation by adding excess lysine. The results indicated that 100 % SurA modification was reached after 15 minutes, suggesting that the activated *o*NBA intermediate is long-lived (**Figure 3D**). The extended lifetime of the *o*NBA intermediate and its stability in aqueous buffer, allows it to retain its specificity and high yields providing sufficient time to profile lysine residues. However, a consequence of increased stability is less sensitive time-resolution when compared to the nanosecond reaction pulse of an activated diazirine species ^29^. *o*NBA groups therefore represent a complementary tool in crosslinker design for time-resolved applications.

**Figure 3.**
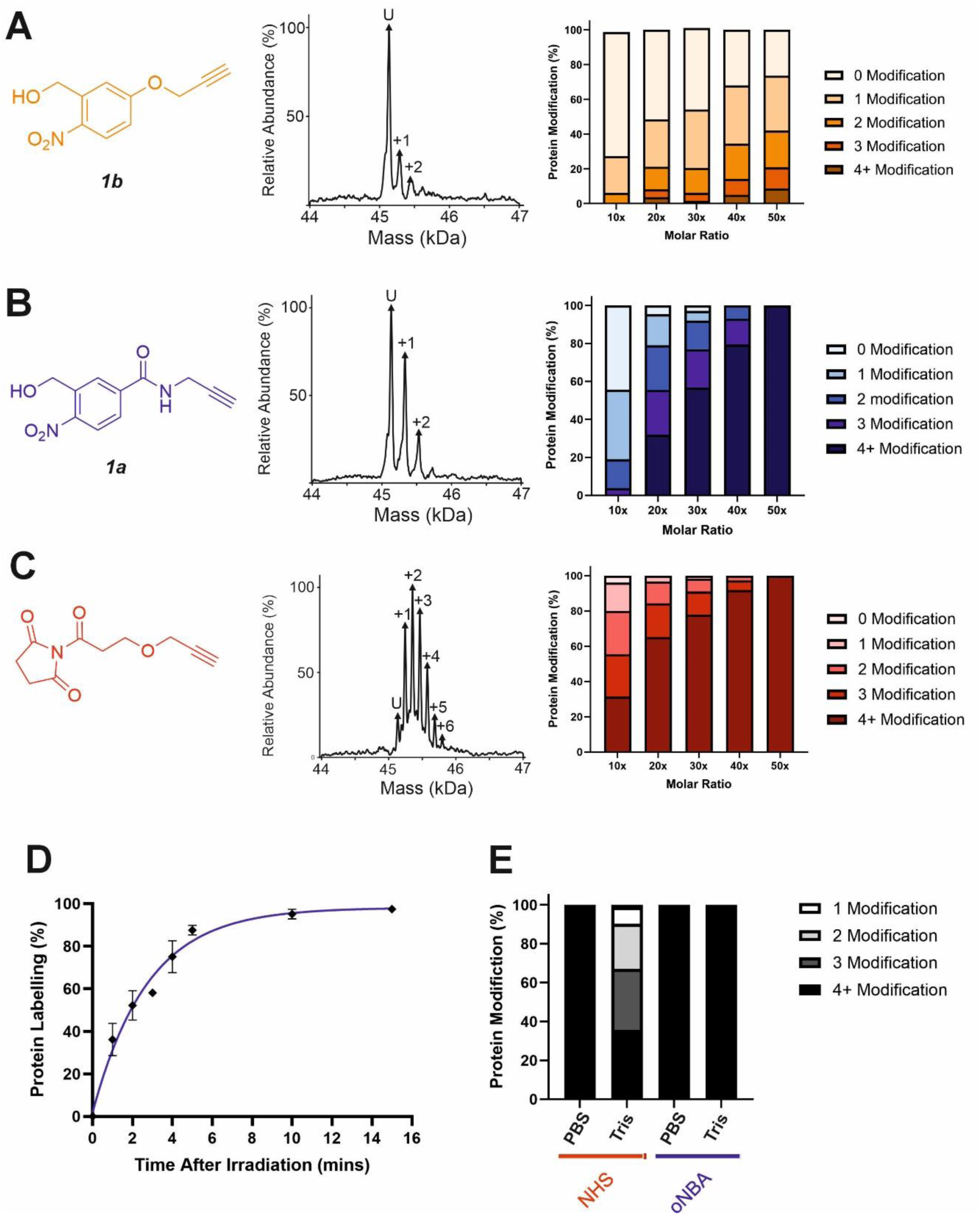
Labelling reactions of a model globular protein, SurA, using amide-oNBA, methoxy oNBA and NHS-ester reactive groups. The labelling efficiency of **(A) 1b; (B) 1a**; and **(C)** NHS-ester reactive groups. All head groups (left) were incubated with SurA at varying molar excesses and samples containing *o*NBA reactive groups were irradiated with 365 nm light for 30 seconds. The total reaction time was 1 hour. An exemplar deconvoluted mass spectra is shown for labelling performed upon addition of the probe at a 1:10 molar excess (centre). The relative abundance of all the species present was quantified using intact MS and the mean values of three replicate measurements are plotted (right). The data used to generate these graphs are detailed in **Table S1, Table S2,** and **Table S3**. **(D)** Labelling reactions were quenched with excess lysine at different time points after irradiation, with labelling performed using *1a*, and the relative abundance of the species present quantified to determine the extent of protein labelling. **(E)** Bar chart highlighting the buffer compatibility of the *o*NBA reactive group *1a* when compared to an NHS-ester group. Labelling reactions were carried out by incubating SurA with a 50-fold molar excess of the head group in PBS or in 20 mM Tris.

A restriction when using NHS-ester based crosslinkers is the need to remove amines from the reaction buffer. This renders the commonly encountered biochemical buffer Tris unsuitable for NHS-ester based XL studies ^30^. In view of this, we chose to perform the previously described labelling reactions in PBS (**Figure 3A - 3D**). To investigate the compatibility of the *o*NBA group for protein-labelling in Tris-based buffers, the labelling reactions were performed in 20 mM Tris-HCl and PBS for both the NHS-ester and **1a** reactive groups. Head group **1a** clearly demonstrated its ability to successfully react with lysine residues even in 20 mM Tris-HCl, whereas the NHS-ester showed a significant decrease in labelling efficiency in this buffer (**Figure 3E**). The improved buffer compatibility of *o*NBA photoproducts could be useful for both labelling reactions and XL-MS in cases where Tris-based buffers are utilised for sample preparation.

### Investigation of *o*NBA-lysine reaction products

Previous work investigating *o*NBA reactivity with lysine residues upon UV-irradiation has highlighted the different potential products formed upon reaction of the ε-amine with the activated *o*NBA species ^31^. The aldehyde formed from the dehydration reaction initiated upon UV irradiation of *o*NBA could allow for the formation of an imine. This product is isobaric to the indazolone product which forms (**Figure 4A**), making the imine and indazolone indistinguishable by MS. Imines are susceptible to acid-induced hydrolysis and reduction, and are therefore likely to decompose under LC-MS/MS conditions, making detection of any imine product by LC-MS/MS unlikely. Another possible product of *o*NBA reaction with lysine is a secondary amine, formed by reduction of the imine (**Figure 4A**), which has been previously reported ^19^. The small difference in mass between these two products (2 Da) is difficult to detect using intact-MS, therefore we utilised high-resolution tandem MS of tryptic peptides to differentiate between the potential products. To achieve this, we labelled bovine serum albumin (BSA) with both *o*NBA analogues, then performed tryptic digestion to detect both modified and unmodified peptides (**Figure S1**). Using label free quantification, we identified the relative abundancies of both indazolone and amine products for both *o*NBA analogues (**Figure 4B**). **1a** exhibited significantly higher labelling yields relative to **1b** and consistently formed the desired indazolone selectively (>20:1 ratio, **Figure 4B**). Reagent **1b** afforded low yields of labelled peptides (<0.5%) and the products comprised roughly equal mixture of indazolone and amine products, consistent with previous literature findings concerning *O*-substituted *o*NBAs ^19^. We propose that the lower reactivity of this group, combined with its lack of specificity, renders this analogue unsuitable for XL-MS (**Figure 4B**). Consequently, the amide-containing *o*NBA group ***1a*** was taken forward and incorporated into our crosslinker design.

**Figure 4.**
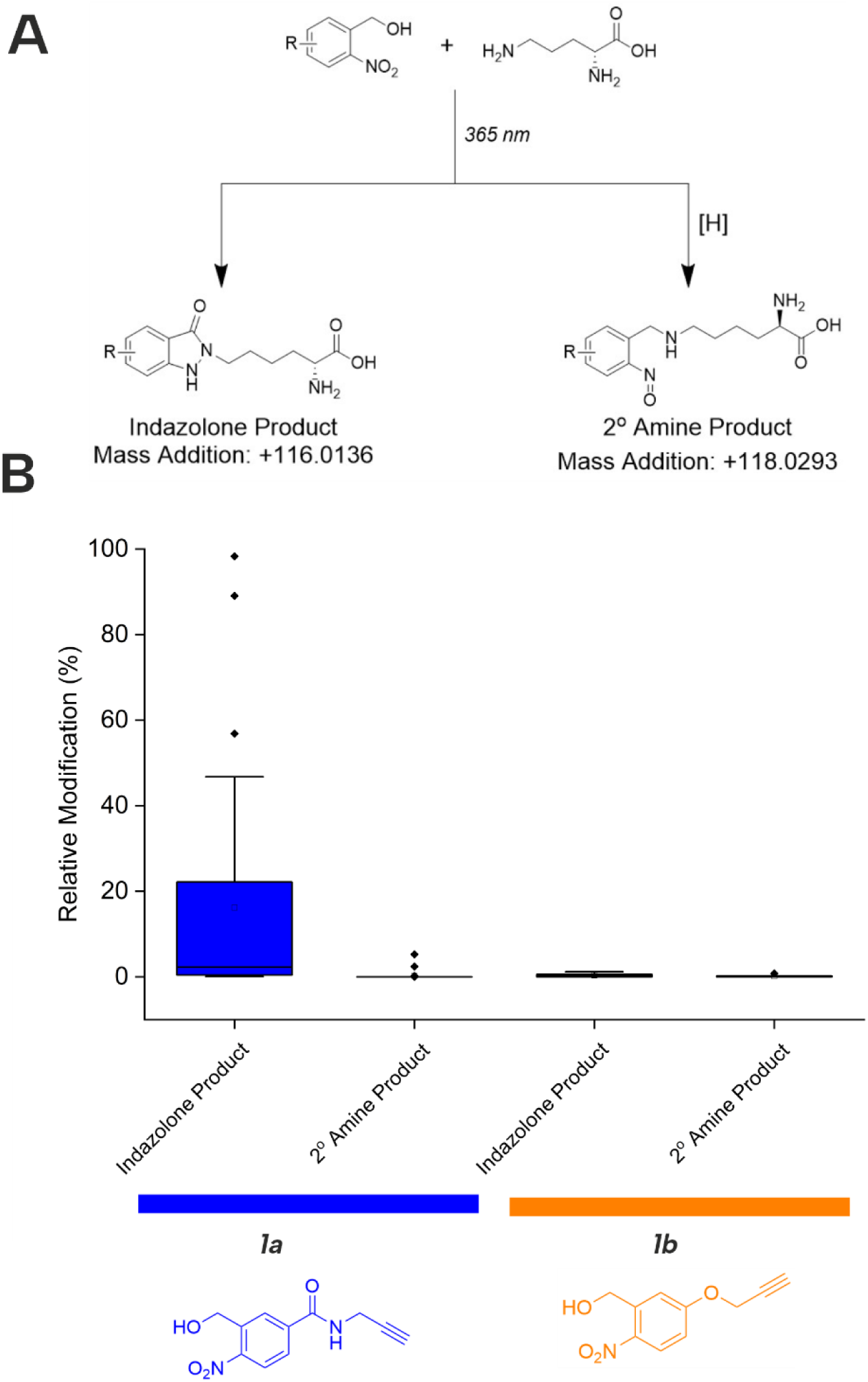
Relative abundance of different oNBA reaction products with lysine residues in BSA quantified by LC-MS. **(A)** Possible products of the oNBA reaction with primary amines detectable by LC-MS, including the monoisotopic mass addition expected from each respective reaction with a lysine sidechain ([H] represents a generic reducing agent). **(B)** Box plots depicting the yield of all modified peptides (relative to their corresponding unmodified counterpart) detected in BSA when labelling was performed with **1a** or **1b**. These data highlight the increased yield of indazolone products using **1a** over **1b**, and the overall greater reaction yields obtained for **1a**. Exemplar MS/MS spectra of modified peptides can be found in **Figure S1**.

### Characterisation of a hetero-bifunctional *o*NBA crosslinker

In analogy with commercial photoactivatable crosslinkers that are hetero-bifunctional, we synthesised a molecule containing NHS-ester and *o*NBA reactive moieties (**3**, **Figure 5A**). NHS-esters are often incorporated into hetero-bifunctional crosslinkers to decorate the protein surface with a reactive group that can then be subsequently activated by UV irradiation ^32^. We applied a similar design principle to generate a crosslinker with the ability for temporally controlled crosslinking via one of the reactive moieties. We crosslinked BSA using **3** and determined successful protein crosslinking using SDS-PAGE, wherein smearing of the gel band corresponding to BSA was observed after crosslinking (**Figure 5A**). The crosslinked samples were digested and analysed by LC-MS/MS. Across the 3 crosslinked samples analysed, 38 total crosslinks were identified with 38% of these found in all samples (**Figure 5B, Figure S2**). The yields of crosslinks were lower than established *bis*-NHS crosslinkers (e.g. DSS)^9^, however, when compared to other hetero-bifunctional photoactivatable crosslinkers (e.g. SDA), the number of detected crosslinks was increased ^33^. We propose that this is likely due to the high specificity of the photochemically activated *o*NBA reactions for lysine sidechains, and the fact that *o*NBA photoproducts are not deactivated by water, unlike diazirines ^34^. The crosslinks were mapped onto the crystal structure of BSA to determine the distance between the residues involved in each crosslink (**Figure 5C**). No identified crosslinks exceeded 40 Å (Cα-Cα Euclidean distance), indicating that this crosslinker provides a viable distance restraint that is consistent with those obtained from other reagents ^35^, making **3** suitable for structural proteomics.

**Figure 5.**
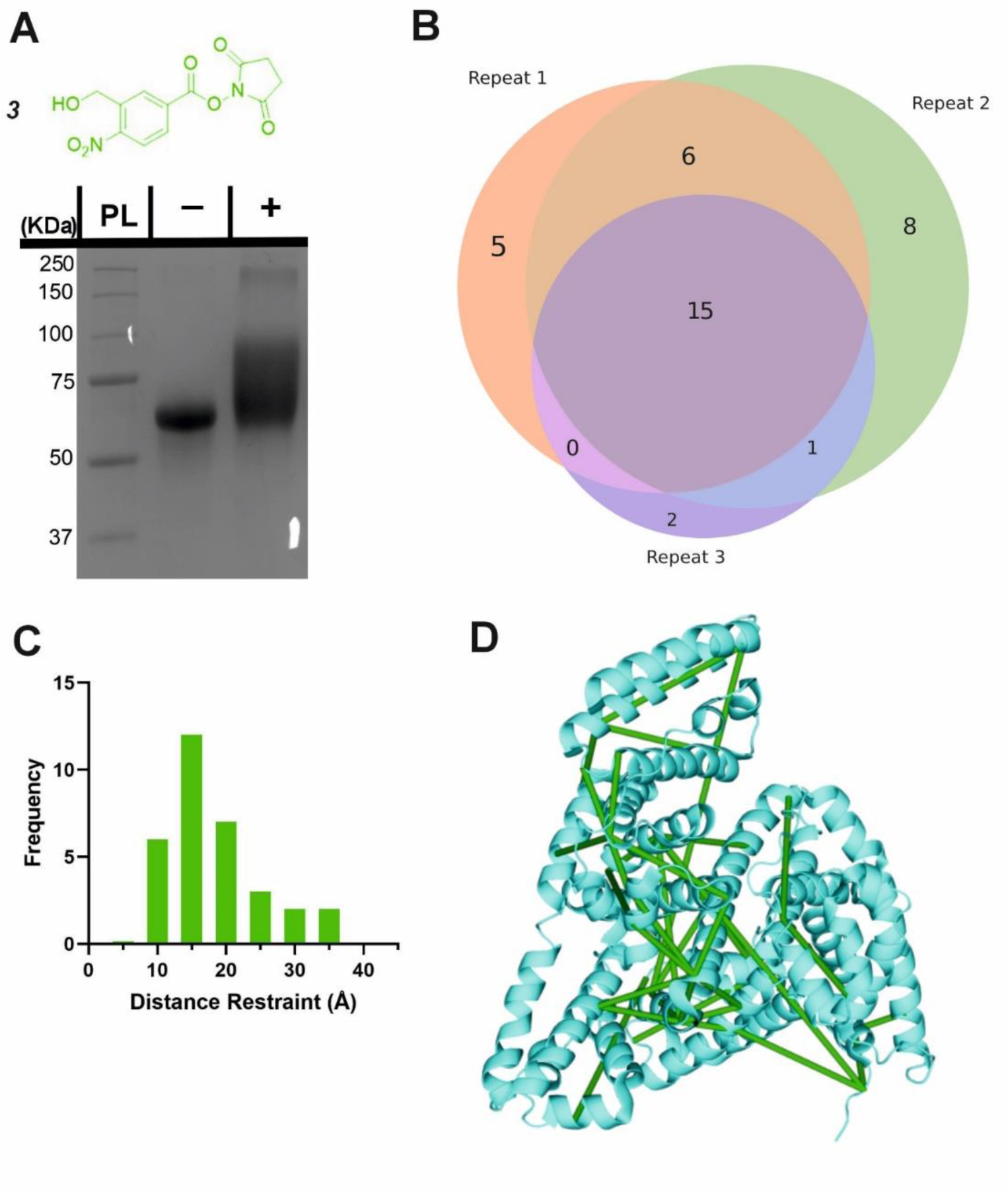
Crosslinking of BSA using the hetero-bifunctional crosslinker 3. **(A)** The structure of the synthesised hetero-bifunctional crosslinker **3** with the SDS-PAGE gel of BSA alone or after reaction with 100-fold molar excess of crosslinker **3**. PL = Protein ladder, + and - indicate that crosslinker was or was not added to the protein, respectively. **(B)** Venn diagram showing the reproducibility of BSA crosslinking using crosslinker **3**. The Venn diagram displays the reproducibility of crosslinks identified in three biological replicates. **(C)** Histogram of the Euclidean distances of crosslinks identified as determined by mapping these onto the BSA crystal structure. **(D)** The crosslinks identified from all 3 replicates mapped onto the BSA crystal structure (PDB: 4F5S) ^36^. Mapping of crosslinks onto structures and distance measurements was performed using the PyXlinkViewer plugin for PyMol^24^. Exemplar MS/MS spectra of crosslinked peptides can be found in **Figure S2**.

### Characterisation of a homo-bifunctional *o*NBA crosslinker

Given the high specificity of *o*NBA chemistry towards ԑ-amines, we aimed to design a crosslinker in a similar style to DSS (which comprises two NHS-esters) that contained two *o*NBA reactive moieties. This new homo-bifunctional crosslinker (**4**, **Figure 6A**) was used to crosslink the model protein BSA. Seventeen total crosslinks were detected by LC-MS/MS (**Figure S3**), fewer than the previously described hetero-bifunctional crosslinker (38 total crosslinks, **Figure 5B**). This was expected as our earlier findings showed that the *o*NBA reactive moiety is less reactive than the NHS-ester (**Figure 2D**). The crosslinks identified were reproducible between biological replicates (**Figure 6B**). These crosslinks were mapped onto a BSA crystal structure and the distances between crosslinked residues calculated (Cα-Cα Euclidean distance), revealing no crosslinks exceeded 40 Å (**Figure 6C**), indicating that this crosslinker provides a viable distance restraint that is consistent with those obtained from other reagents ^35^, making **4** suitable for structural proteomics.

**Figure 6.**
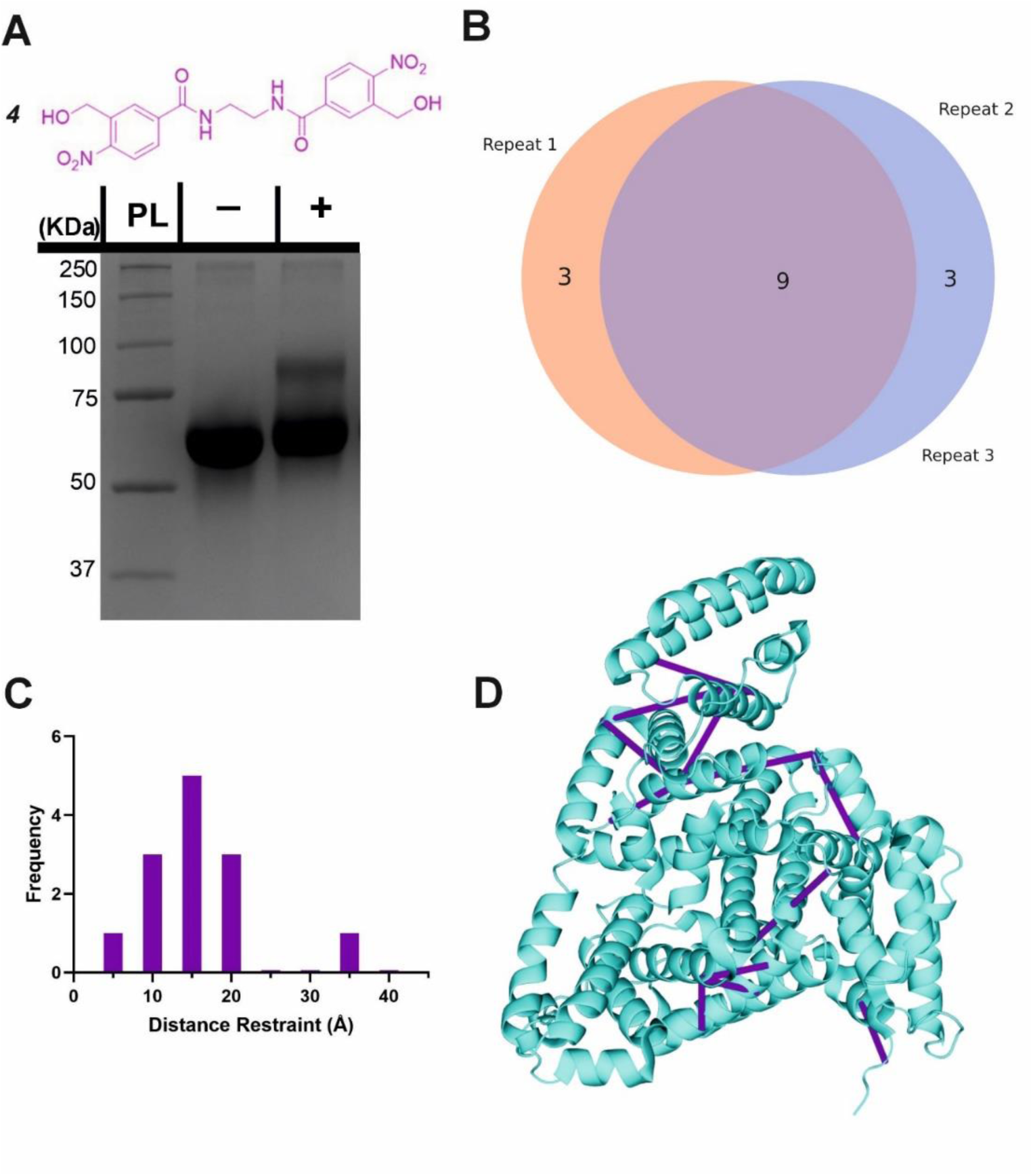
Crosslinking of BSA using the homo-bifunctional crosslinker 4. **(A)** The structure of the synthesised homo-bifunctional crosslinker **4** with the SDS-PAGE gel of BSA alone or after reaction with 100-fold molar excess of crosslinker **4**. PL = Protein ladder, + and - indicate that crosslinker was or was not added to the protein, respectively. **(B)** Venn diagram showing the reproducibility of crosslinking using crosslinker **4**. The Venn diagram displays the reproducibility of crosslinks identified in three biological replicates. **(C)** Histogram of the Euclidean distances of crosslinks identified as determined by mapping these onto the BSA crystal structure. **(D)** The crosslinks identified from all 3 replicates mapped onto the crystal structure (PDB: 4F5S) ^36^. Mapping of crosslinks onto structures and distance measurements was performed using the PyXlinkViewer plugin for PyMol ^24^. Exemplar MS/MS spectra of crosslinked peptides can be found in **Figure S3**.

### Characterising Protein-Protein Interaction using Novel Crosslinkers

After successfully identifying crosslinks generated by both hetero-bifunctional and homo-bifunctional *o*NBA crosslinkers, we next sought to characterise a protein-protein interaction (PPI), using the SurA-OmpX complex as a model system to identify interprotein crosslinks. These proteins are known binding partners and previous crosslinking data exists using both NHS ester and diazirine-based reagents ^21^. To exploit the heterobifunctional nature of **3**, which contains both a constitutively active NHS-ester and a photoactivatable *o*NBA group, we incubated SurA with **3** for 30 minutes to enable the NHS-ester to react with lysine residues. After this, we introduced OmpX to form the SurA-OmpX complex. The sample was then irradiated with UV light to generate interprotein crosslinks between SurA and OmpX. This strategy represents a ‘plant and cast’ approach to crosslinking, which has been previously exploited by others^37^. Using crosslinker **3**, we identified 79 unique crosslinks including 9 interprotein crosslinks (**Figure 7A**, **Figure S4**). All the interprotein crosslinks between SurA and OmpX were localised onto K60 of OmpX. This is consistent with previously published XL-MS data of this complex, where a diffuse interaction interface was captured, and highlights the ability of crosslinker **3** to describe protein binding interfaces in heteromeric assemblies ^21^.

**Figure 7.**
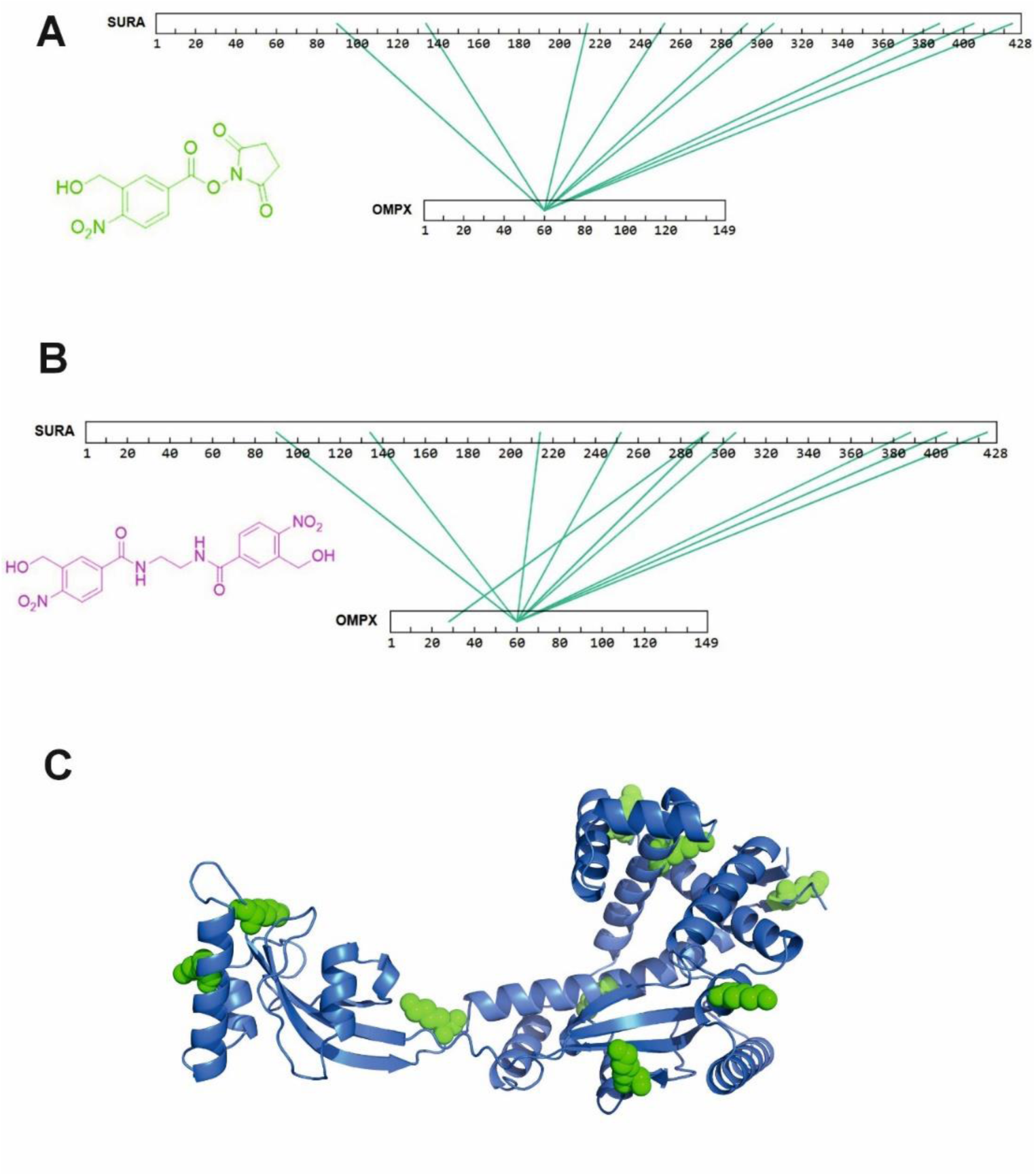
XL-MS analysis of SurA and OmpX with synthesised hetero-bifunctional and homo-bifunctional *o*NBA-based crosslinkers. (A) SurA and OmpX were crosslinked using 100-fold molar excess of the synthesised crosslinker **3**. The interprotein crosslinks identified were mapped onto the protein sequences using XiView ^38^. **(B)** SurA and OmpX were crosslinked using 100-fold molar excess of crosslinker **4**. The interprotein crosslinks identified were mapped onto the protein sequences using XiView. (C) The crystal structure of SurA (PDB: 1M5Y) ^39^. The green spheres represent the 9 lysine residues on SurA that formed crosslinks with K60 of OmpX. Exemplar MS/MS spectra of crosslinked peptides can be found in **Figure S4** and **Figure S5**.

As crosslinker **4** comprises two *o*NBA moieties which are both UV-activatable, a ‘plant and cast’ strategy could not be utilised for this reagent. Instead, we formed the SurA-OmpX complex and then added crosslinker **4**, activating its two end groups simultaneously via UV irradiation. We identified 58 crosslinks with 10 interprotein crosslinks (**Figure S5**). The crosslinks were again mainly localised onto K60 of OmpX (**Figure 7B**). The 9 interprotein crosslinks identified using crosslinker **3** were also present when using crosslinker **4**, suggesting that comparable data can be generated using the two different crosslinkers. The lysine residues of SurA that forms crosslinks with K60 of OmpX were mapped onto the crystal structure of SurA (**Figure 7C**). This highlights the large dispersion of lysine residues across the sequence of SurA and agrees with previous work that has used XL-MS to demonstrate that these two binding partners have a diffuse interaction interface. The consistency of our findings relative to previous XL-MS studies and other complimentary techniques ^21^ further supports the potential of *o*NBA crosslinkers in unveiling the architectures of protein complexes.

### Investigating ‘mono-link’ products when using an *o*NBA based crosslinker

The propensity of crosslinking reagents to form uninformative mono-links instead of the desired crosslinks is a common challenge in XL-MS, especially for NHS-ester based reagents ^10^. This is due to the susceptibility of the reactive groups used in crosslinkers to irreversible hydrolysis. While the reactive intermediate formed by *o*NBA photolysis also reacts with water, this process is reversible ^40^. After demonstrating the ability of crosslinkers **3** and **4** to detect PPIs, we investigated their propensity to form mono-links. Using database searches, we sought to identify the structures of potential mono-link products formed by both crosslinkers from the SurA-OmpX datasets (**Figure 8A/B**).

**Figure 8.**
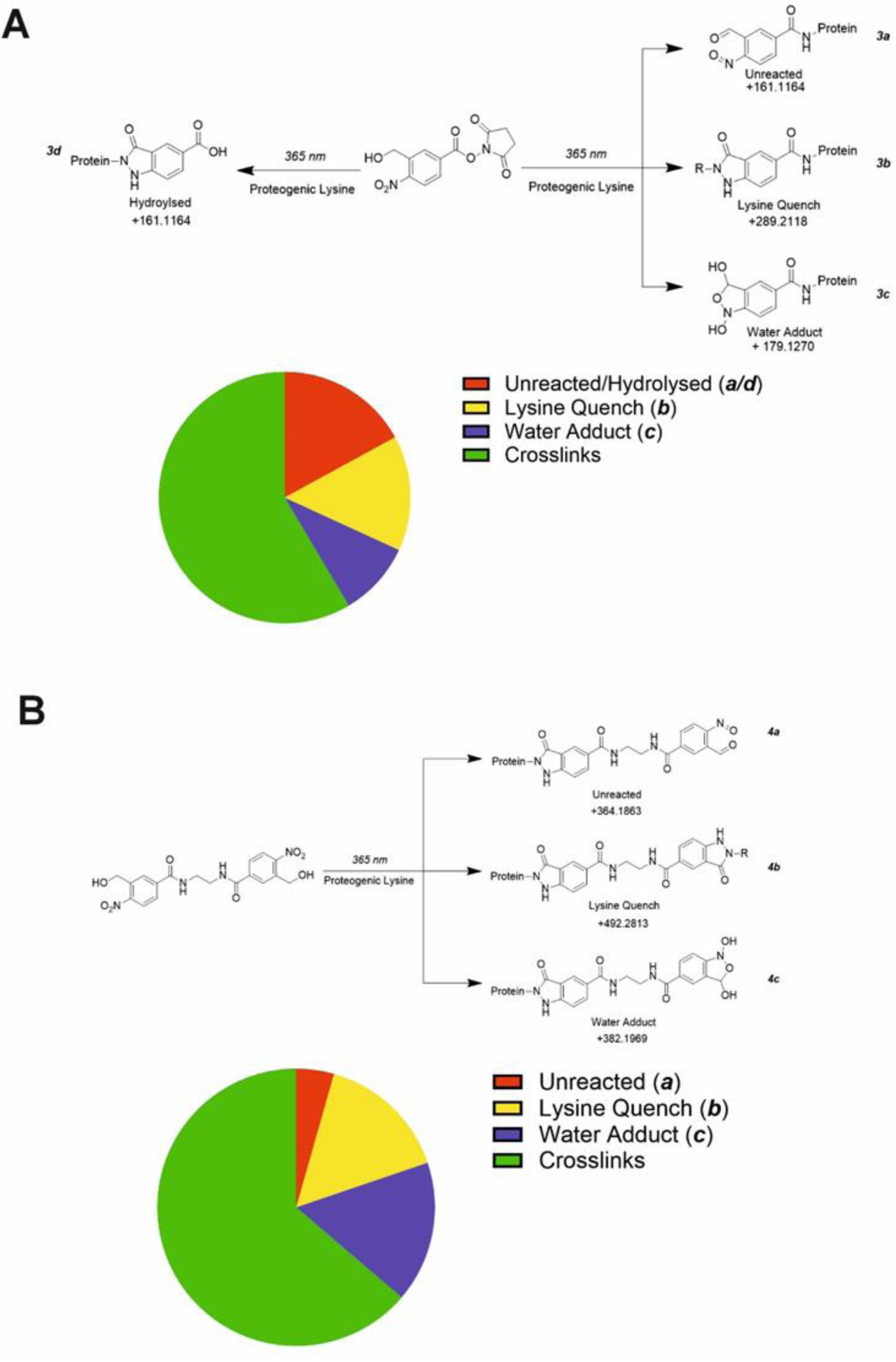
The mono-link products detected using both crosslinkers (3) and (4). **(A)** MS/MS data from the SurA-OmpX crosslinking dataset was searched for potential mono-link products using crosslinker ***3***. The hypothesised mono-links that could be present in the sample were assumed to result from either water or lysine quenching/reactions. The pie chart represents the number of identified unique crosslink/mono-link species within the sample. **(B)** MS/MS data from the SurA-OmpX crosslinking dataset was searched for potential mono-link products using crosslinker ***4***. The hypothesised mono-links that could be present in the sample were assumed to result from either water or lysine quenching/reactions. The pie chart represents the number of identified unique crosslink/mono-link species within the sample.

Using crosslinker **3**, we hypothesised 4 potential dead-end products might form (**3a**, **3b**, **3c**, **3d**, **Figure 8A**). The protein modifications corresponding to irreversible hydrolysis of NHS-ester (**3d**) and the activated oNBA remaining unreacted (**3a**) are isobaric and indistinguishable (**Figure 8A**). Therefore, these species are grouped together in this analysis. We identified 20 **3b** products, 13 **3c** products and 23 **3a/d** products (**Figure 8A, Figure S6**). There was no consistent mono-link product observed when using crosslinker **3**, likely due to the diversity of products available. The number of potential mono-link products is reduced to three when using crosslinker **4**, as it is homo-bifunctional. The potential mono-links identified comprised 14 lysine quenched products (**4b**), 15 water adducts (**4c**) and 4 unreacted oNBA moieties (**4a**) (**Figure 8B, Figure S7**). The reduced number of mono-link identifications when using crosslinker **4** compared to crosslinker **3**, suggests that unreacted *o*NBA moieties are not commonly present and the high identification of such structures when using crosslinker **3** is likely due to the isobaric irreversible hydrolysis of the NHS-ester group (**3d**).

Overall, we identified 56 and 33 unique modifications for crosslinker **3** and ***4*** respectively. The percentage of unique crosslinks identified when compared to the number of unique mono-links was 66 % for **3** and 70 % for **4**, meaning the number of identified crosslinks is *ca.* twice the number of identified mono-links (**Figure 8A,B**). The high number of crosslinks compared to mono-links seen here differs from previous work utilising NHS-ester based reagents, which often have a higher propensity to form mono-links over crosslinks ^41^. This represents a significant advantage over existing crosslinking groups and is potentially due to the high specificity and reactivity of the *o*NBA group for primary amines, in addition to its reversible reaction with water, reducing the likelihood of side-reactions that are the cause of mono-link formation.

## Conclusion

Here we have developed new *o*NBA reactive warheads and designed novel chemical crosslinkers for mapping protein-protein interactions. By synthesising and characterising two *o*NBA reactive head groups, we highlight the importance of substitution pattern in determining the behaviour of *o*NBA chemistry for biological applications involving UV-irradiation/activation. We have demonstrated that once appropriately understood, UV photoactivation of *o*NBA can generate high yields of modified protein with highly predictable reaction products, demonstrating that *o*NBA based reagents are suitable for protein labelling. Two *o*NBA based crosslinkers, a hetero-bifunctional and homo-bifunctional reagent, were synthesised and both reliably formed crosslinks detectable by mass spectrometry. These reagents can be used to generate distance restraints that are comparable to those that can be obtained with most commercially available NHS-ester based crosslinkers, demonstrating the possible applications of these reagents in structural proteomics. The lower propensity to form mono-links when using *o*NBA based crosslinkers to measure interprotein interactions also addresses the limitations of traditional photoactivatable crosslinkers, expanding the scope of XL-MS. An additional benefit of the *o*NBA reactivity profile is the high specificity towards lysine residues. This simplifies the search space for crosslinking data analysis and may prove extremely beneficial when working with more complex samples. We anticipate that the selectivity and bio-orthogonal activation of *o*NBA crosslinking reagents will further contribute to the future understanding of protein structure and interactions, including in dynamic systems, by exploiting the temporal control afforded by these novel reagents.

## Acknowledgements

We thank the members of the Calabrese, Wright, Radford and Brockwell groups for helpful discussion, along with Gemma Wildsmith for technical support and advice within the mass spectrometry facility. A.K.C acknowledges support from an EPSRC studentship (EP/W524372/1 project reference 2879844). M.W. and A.H. acknowledge funding from BBSRC (BB/Y00034X/1, BB/V003577/2). K.L. acknowledges funding and support from the School of Chemistry, University of Leeds. A.N.C. acknowledges support of a Sir Henry Dale Fellowship jointly funded by the Wellcome Trust and the Royal Society (220628/Z/20/Z). S.E.R. acknowledges funding from the Royal Society (RSRP\R1\211057). Funding from Wellcome (223810/Z/21/Z) and NIHR (NIHR200633) enabled the purchase of mass spectrometry equipment.

## Data Availability

The mass spectrometry proteomics data have been deposited to the ProteomeXchange Consortium via the PRIDE ^42^ partner repository with the dataset identifier PXD063033.

## Author Contributions

**Adam Cahill**: Methodology, Software, Validation, Formal analysis, Investigation, Data curation, Writing – original draft, Visualization. **Martin Walko**: Methodology, Validation, Resources, Writing – Review and Editing, Supervision. **Benjamin Fenton**: Resources, Writing & Editing. **Sri Ranjani Ganji**: Methodology, Resources, Writing – Review & Editing**. Anne Herbert**: Resources, Writing – Review & Editing. **Sheena E. Radford**: Supervision, Project administration, Funding acquisition. **Nikil Kapur**: Methodology, Resources, Writing – Review & Editing. **Keith Livingstone**: Writing – Review & Editing, Supervision, Project administration. **Megan H. Wright**: Conceptualization, methodology, Validation, Writing – Review & Editing, Supervision, Funding acquisition. **Antonio N. Calabrese**: Conceptualization, Methodology, Validation, Writing – Review & Editing, Visualisation, Supervision, Project administration, Funding acquisition.

## References

1 K. Stahl, R. Warneke, L. Demann, R. Bremenkamp, B. Hormes, O. Brock, J. Stülke and J. Rappsilber, Nat Commun, 2024, 15, 7866.

2 J. Luo and J. Ranish, Elife, 2024, 13, RP99809, DOI:10.7554/eLife.99809.

3 J. W. Back, M. A. Sanz, L. de Jong, L. J. de Koning, L. G. J. Nijtmans, C. G. de Koster, L. A. Grivell, H. van der Spek and A. O. Muijsers, Protein Science, 2002, 11, 2471–2478.

4 B. Yang, H. Wu, P. D. Schnier, Y. Liu, J. Liu, N. Wang, W. F. DeGrado and L. Wang, Proceedings of the National Academy of Sciences, 2018, 115, 11162–11167.

5 H. Gao, Q. Zhao, Z. Gong, B. Zhong, J. Chen, Z. Sui, X. Li, Z. Liang, Y. Zhang and L. Zhang, Anal Chem, 2022, 94, 12398–12406.

6 C. Gutierrez, L. J. Salituro, C. Yu, X. Wang, S. F. DePeter, S. D. Rychnovsky and L. Huang, Molecular & Cellular Proteomics, 2021, 20, 100084.

7 R. Huang, W. Zhu, Y. Wu, J. Chen, J. Yu, B. Jiang, H. Chen and W. Chen, Chem Sci, 2019, 10, 6443–6447.

8 A. Kao, C. Chiu, D. Vellucci, Y. Yang, V. R. Patel, S. Guan, A. Randall, P. Baldi, S. D. Rychnovsky and L. Huang, Molecular & Cellular Proteomics, 2011, 10, M110.002170.

9 C. Iacobucci, C. Piotrowski, R. Aebersold, B. C. Amaral, P. Andrews, K. Bernfur, C. Borchers, N. I. Brodie, J. E. Bruce, Y. Cao, S. Chaignepain, J. D. Chavez, S. Claverol, J. Cox, T. Davis, G. Degliesposti, M.-Q. Dong, N. Edinger, C. Emanuelsson, M. Gay, M. Götze, F. Gomes-Neto, F. C. Gozzo, C. Gutierrez, C. Haupt, A. J. R. Heck, F. Herzog, L. Huang, M. R. Hoopmann, N. Kalisman, O. Klykov, Z. Kukačka, F. Liu, M. J. MacCoss, K. Mechtler, R. Mesika, R. L. Moritz, N. Nagaraj, V. Nesati, A. G. C. Neves-Ferreira, R. Ninnis, P. Novák, F. J. O’Reilly, M. Pelzing, E. Petrotchenko, L. Piersimoni, M. Plasencia, T. Pukala, K. D. Rand, J. Rappsilber, D. Reichmann, C. Sailer, C. P. Sarnowski, R. A. Scheltema, C. Schmidt, D. C. Schriemer, Y. Shi, J. M. Skehel, M. Slavin, F. Sobott, V. Solis-Mezarino, H. Stephanowitz, F. Stengel, C. E. Stieger, E. Trabjerg, M. Trnka, M. Vilaseca, R. Viner, Y. Xiang, S. Yilmaz, A. Zelter, D. Ziemianowicz, A. Leitner and A. Sinz, Anal Chem, 2019, 91, 6953–6961.

10 M. Sinnott, S. Malhotra, M. S. Madhusudhan, K. Thalassinos and M. Topf, Structure, 2020, 28, 1061–1070.e3.

11 A. F. Gomes and F. C. Gozzo, Journal of Mass Spectrometry, 2010, 45, 892–899.

12 R. K. Venkatraman and A. J. Orr-Ewing, J Am Chem Soc, 2019, 141, 15222–15229.

13 D. A. Modarelli, S. Morgan and M. S. Platz, J Am Chem Soc, 1992, 114, 7034–7041.

14 A. V. West, G. Muncipinto, H.-Y. Wu, A. C. Huang, M. T. Labenski, L. H. Jones and C. M. Woo, J Am Chem Soc, 2021, 143, 6691–6700.

15 C. Wang, Y. Liu, C. Bao, Y. Xue, Y. Zhou, D. Zhang, Q. Lin and L. Zhu, Chemical Communications, 2020, 56, 2264–2267.

16 A.-D. Guo, K.-H. Wu and X.-H. Chen, RSC Adv, 2021, 11, 2235–2241.

17 A.-D. Guo, D. Wei, H.-J. Nie, H. Hu, C. Peng, S.-T. Li, K.-N. Yan, B.-S. Zhou, L. Feng, C. Fang, M. Tan, R. Huang and X.-H. Chen, Nat Commun, 2020, 11, 5472.

18 Y. Xu, H. Hu, Y. Ran, W. Zhao, A.-D. Guo, H.-J. Nie, L. Zhai, G.-L. Yin, J.-T. Cheng, S. Tao, B. Yang, M. Tan and X.-H. Chen, 2025, BiorXiv, preprint available, DOI: 10.1101/2025.03.18.643847.

19 X. Chen, C. Jiménez López, A. Nadler and F. Stengel, ChemBioChem, 2024, 25, e202400620.

20 A. Romano, I. Roppolo, M. Giebler, K. Dietliker, Š. Možina, P. Šket, I. Mühlbacher, S. Schlögl and M. Sangermano, RSC Adv, 2018, 8, 41904–41914.

21 A. N. Calabrese, B. Schiffrin, M. Watson, T. K. Karamanos, M. Walko, J. R. Humes, J. E. Horne, P. White, A. J. Wilson, A. C. Kalli, R. Tuma, A. E. Ashcroft, D. J. Brockwell and S. E. Radford, Nat Commun, 2020, 11, 2155.

22 M. T. Marty, A. J. Baldwin, E. G. Marklund, G. K. A. Hochberg, J. L. P. Benesch and C. V. Robinson, Anal Chem, 2015, 87, 4370–4376.

23 C. Iacobucci, M. Götze, C. H. Ihling, C. Piotrowski, C. Arlt, M. Schäfer, C. Hage, R. Schmidt and A. Sinz, Nat Protoc, 2018, 13, 2864–2889.

24 B. Schiffrin, S. E. Radford, D. J. Brockwell and A. N. Calabrese, Protein Science, 2020, 29, 1851– 1857.

25 J. Brunner, H. Senn and F. M. Richards, J Biol Chem, 1980, 255, 3313–8.

26 A. Belsom, G. Mudd, S. Giese, M. Auer and J. Rappsilber, Anal Chem, 2017, 89, 5319–5324.

27 A. L. Santos, V. Oliveira, I. Baptista, I. Henriques, N. C. M. Gomes, A. Almeida, A. Correia and Â. Cunha, Arch Microbiol, 2013, 195, 63–74.

28 J. E. Horne, M. Walko, A. N. Calabrese, M. A. Levenstein, D. J. Brockwell, N. Kapur, A. J. Wilson and S. E. Radford, Angewandte Chemie International Edition, 2018, 57, 16688–16692.

29 D. A. Modarelli, S. Morgan and M. S. Platz, J Am Chem Soc, 1992, 114, 7034–7041.

30 E. D. Merkley, J. R. Cort and J. N. Adkins, J Struct Funct Genomics, 2013, 14, 77–90.

31 A.-D. Guo, D. Wei, H.-J. Nie, H. Hu, C. Peng, S.-T. Li, K.-N. Yan, B.-S. Zhou, L. Feng, C. Fang, M. Tan, R. Huang and X.-H. Chen, Nat Commun, 2020, 11, 5472.

32 A. Belsom and J. Rappsilber, Curr Opin Chem Biol, 2021, 60, 39–46.

33 Y. Jiang, X. Zhang, H. Nie, J. Fan, S. Di, H. Fu, X. Zhang, L. Wang and C. Tang, Nat Commun, 2024, 15, 6060.

34 Y. Jiang, X. Zhang, H. Nie, J. Fan, S. Di, H. Fu, X. Zhang, L. Wang and C. Tang, Nat Commun, 2024, 15, 6060.

35 E. D. Merkley, S. Rysavy, A. Kahraman, R. P. Hafen, V. Daggett and J. N. Adkins, Protein Sci, 2014, 23, 747–59.

36 A. Bujacz, Acta Crystallogr D Biol Crystallogr, 2012, 68, 1278–1289.

37 B. Yang, H. Wu, P. D. Schnier, Y. Liu, J. Liu, N. Wang, W. F. DeGrado and L. Wang, Proceedings of the National Academy of Sciences, 2018, 115, 11162–11167.

38 C. W. Combe, M. Graham, L. Kolbowski, L. Fischer and J. Rappsilber, J Mol Biol, 2024, 436, 168656.

39 E. Bitto and D. B. McKay, Structure, 2002, 10, 1489–1498.

40 A. Romano, I. Roppolo, E. Rossegger, S. Schlögl and M. Sangermano, Materials, 2020, 13, 2777.

41 B. Steigenberger, R. J. Pieters, A. J. R. Heck and R. A. Scheltema, ACS Cent Sci, 2019, 5, 1514– 1522.

42 Perez-Riverol Y, Bandla C, Kundu DJ, Kamatchinathan S, Bai J, Hewapathirana S, John NS, Prakash A, Walzer M, Wang S, Vizcaíno JA. Nucleic Acids Res, 2025 6; D543–D553.

