## Supplementary Information for "Design and Characterisation of Photoactivatable and Lysine Reactive *o*-Nitrobenzyl Alcohol-Based Crosslinkers"

### NMR Spectra of Synthesised Compounds

#### 4-(hydroxymethyl)-3-nitro-N-2-propyn-1-ylbenzamide (1a)

$^{1}$H NMR (500 MHz, DMSO-$d_6$)

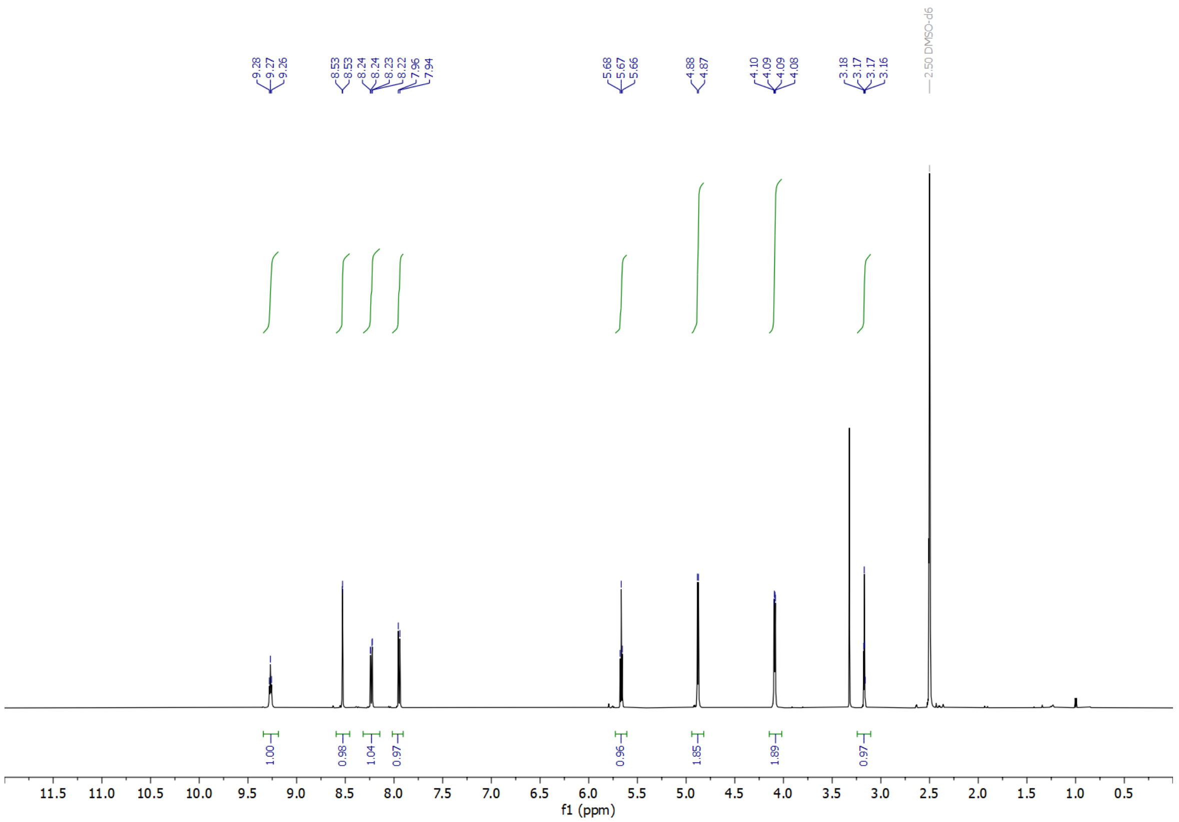

$^{13}$C NMR (101 MHz, DMSO-$d_6$)

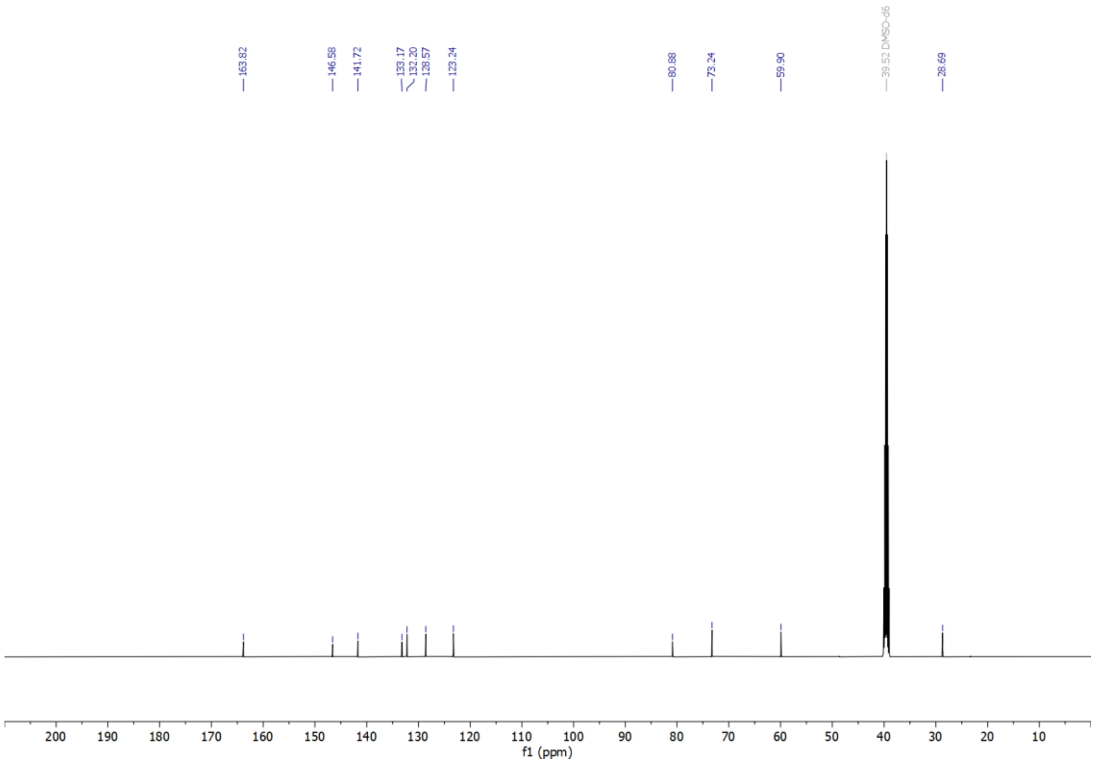

**(2-Nitro-5-(prop-2-yn-1-yloxy)phenyl)methanol (1b)**

$^{1}$H NMR (500 MHz, DMSO-$d_6$)

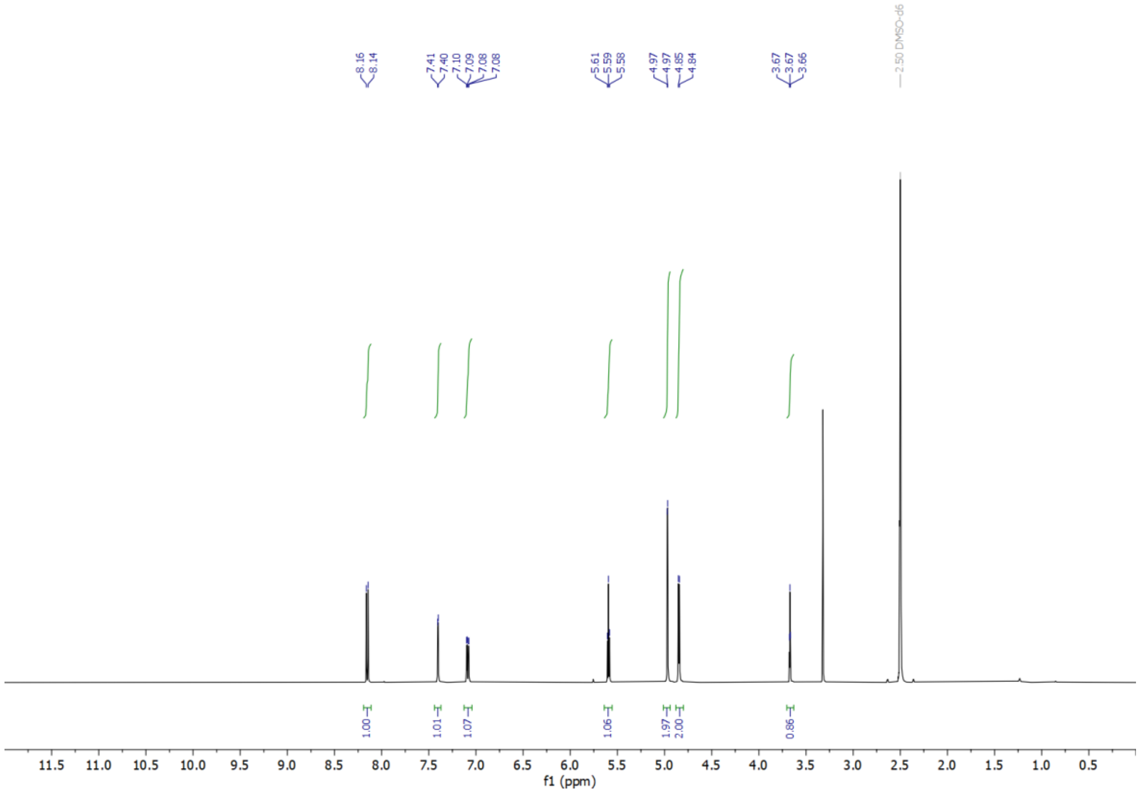


$^{13}$C NMR (101 MHz, DMSO-$d_6$)

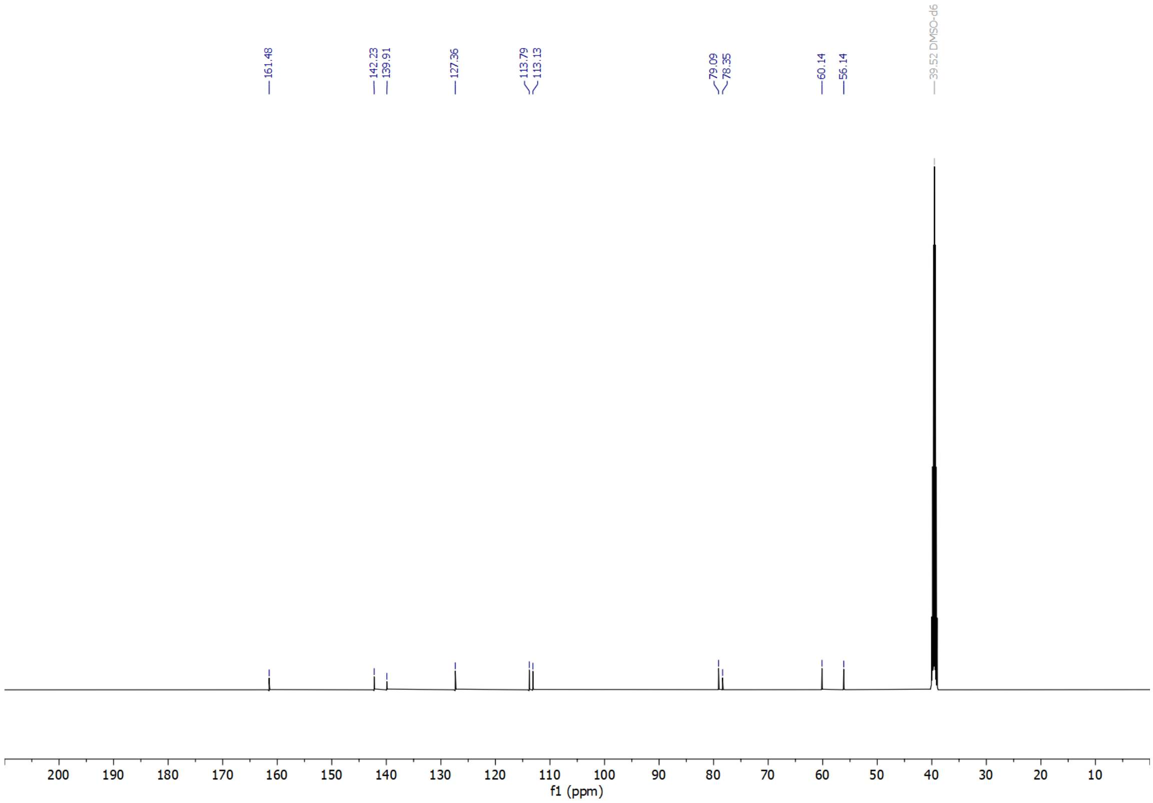

**2,5-dioxo-1-pyrrolidinyl 4-(hydroxymethyl)-3-nitrobenzoate (2)**

$^{1}H$ NMR (500 MHz, DMSO-$d_6$)

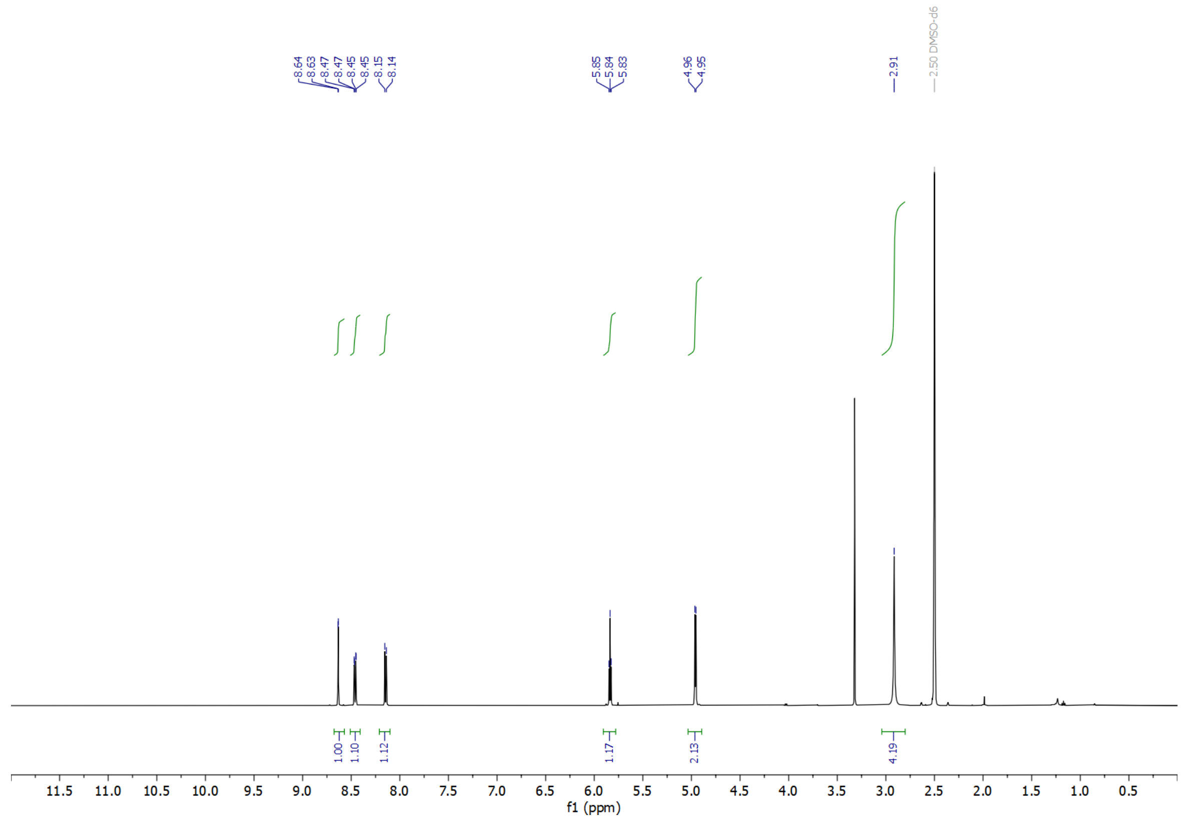


$^{13}C$ NMR (101 MHz, DMSO-$d_6$)

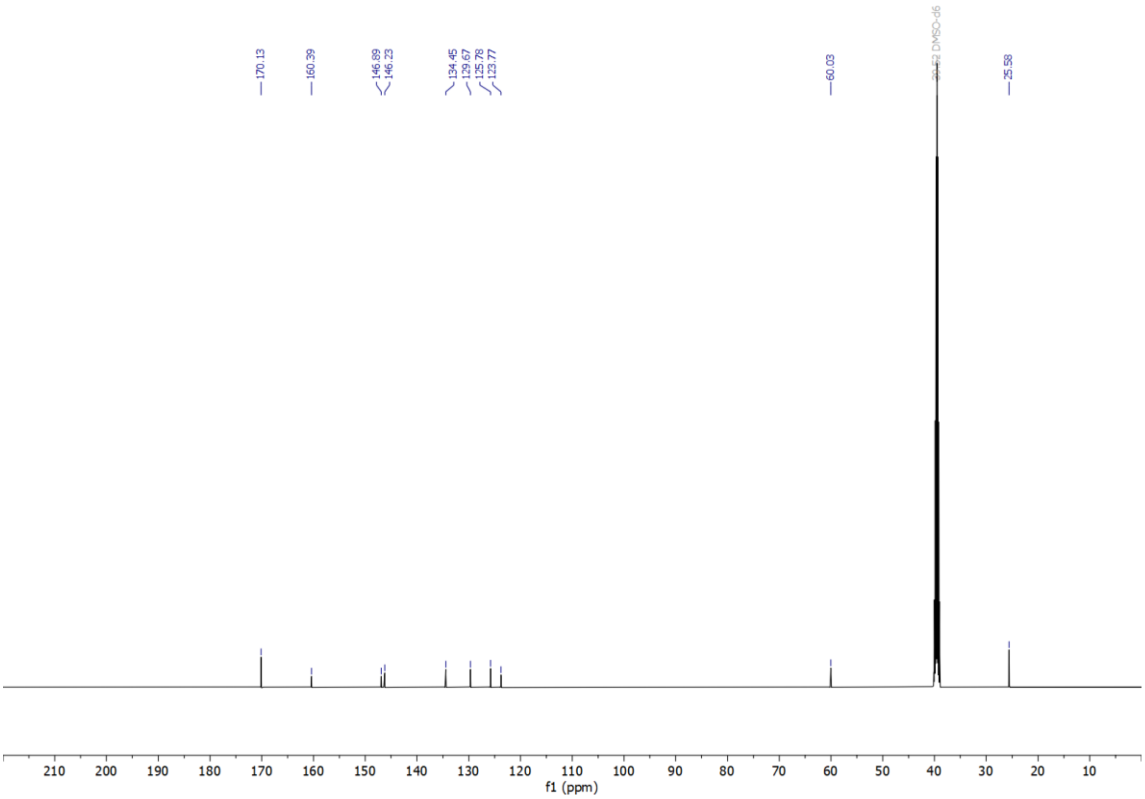

**N,N'-1,2-ethanediylbis(3-hydroxymethyl-4-nitrobenzamide (3)**

$^{1}$H NMR (500 MHz, DMSO-$d_6$)

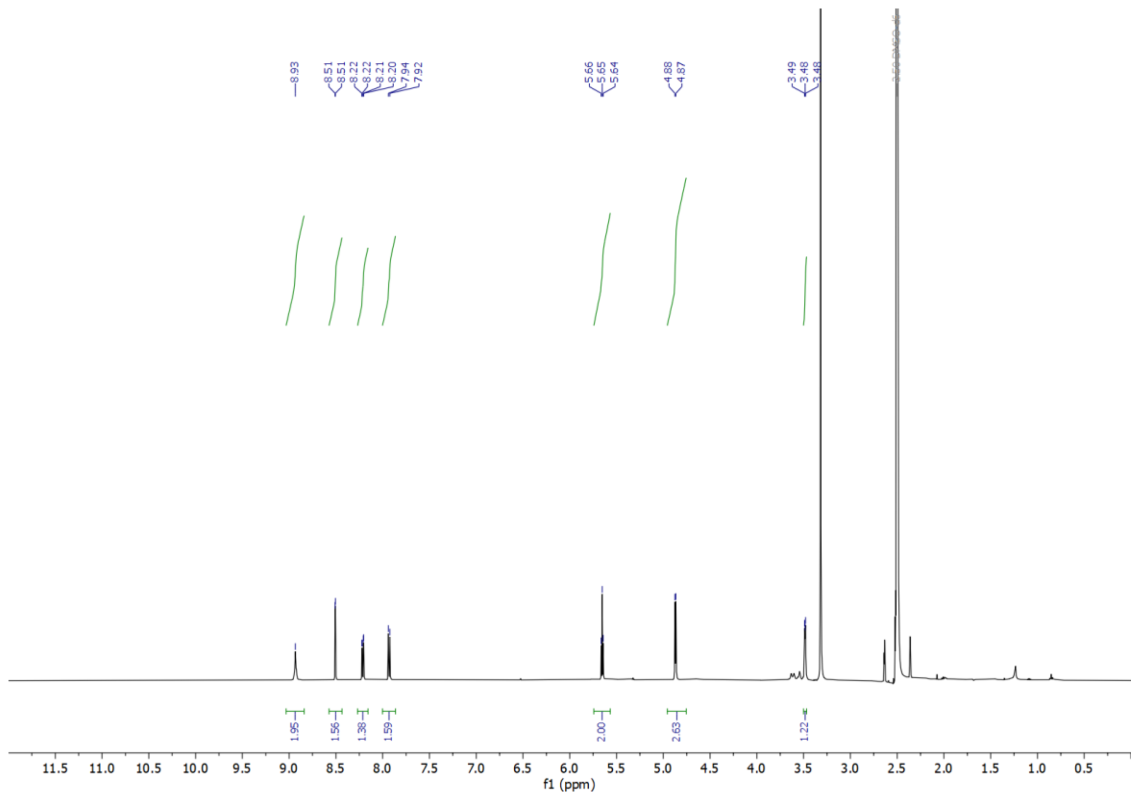


$^{13}$C NMR (101 MHz, DMSO-$d_6$)

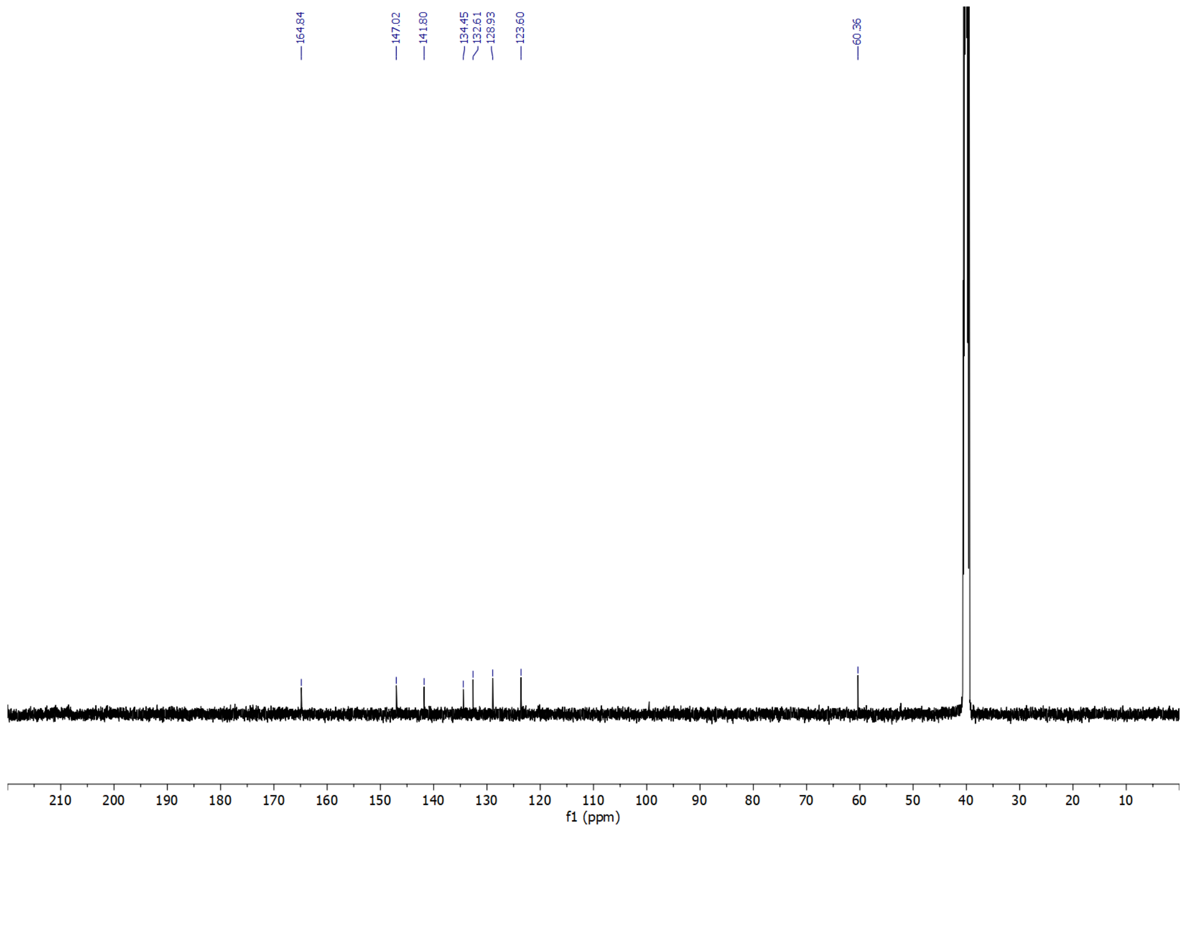

**Supplementary Table 1. Extent of labelling of SurA using *1b* determined from intact MS when the probe was added at different molar excesses.** Data are shown as mean and standard deviation of three replicate measurements.

| | 10x | | 20x | | 30x | | 40x | | 50x | |
|---|---|---|---|---|---|---|---|---|---|---|
| **Modifications** | **Mean** | **SD** | **Mean** | **SD** | **Mean** | **SD** | **Mean** | **SD** | **Mean** | **SD** |
| **0** | 71.377 | 3.929 | 51.558 | 9.415 | 46.880 | 1.877 | 32.062 | 2.220 | 26.450 | 8.440 |
| **1** | 21.181 | 2.017 | 27.313 | 1.380 | 33.645 | 1.589 | 33.489 | 1.054 | 31.583 | 1.598 |
| **2** | 6.080 | 1.331 | 12.939 | 1.012 | 14.283 | 2.353 | 20.362 | 0.738 | 21.165 | 2.068 |
| **3** | 0 | 0 | 4.564 | 4.127 | 4.655 | 4.072 | 9.081 | 1.109 | 12.194 | 1.177 |
| **4+** | 0 | 0 | 3.626 | 3.157 | 1.516 | 2.626 | 5.006 | 0.652 | 8.608 | 8.898 |

**Supplementary Table 2. Extent of labelling of SurA using *1a* determined from intact MS when the probe was added at different molar excesses.** Data are shown as mean and standard deviation of three replicate measurements.

| | 10x | | 20x | | 30x | | 40x | | 50x | |
|---|---|---|---|---|---|---|---|---|---|---|
| Modifications | Mean | SD | Mean | SD | Mean | SD | Mean | SD | Mean | SD |
| 0 | 44.419 | 3.456 | 4.586 | 2.009 | 2.853 | 3.043 | 0 | 0 | 0 | 0 |
| 1 | 36.615 | 1.348 | 16.271 | 2.505 | 5.082 | 1.823 | 0 | 0 | 0 | 0 |
| 2 | 15.069 | 2.186 | 23.643 | 0.805 | 14.981 | 3.124 | 6.946 | 1.701 | 0 | 0 |
| 3 | 3.897 | 0.952 | 23.538 | 1.057 | 20.176 | 2.830 | 13.616 | 2.593 | 0 | 0 |
| 4+ | 0 | 0 | 31.962 | 4.448 | 56.908 | 10.535 | 79.439 | 3.494 | 100 | 0 |

**Supplementary Table 3. Extent of labelling of SurA using *the* NHS-ester probe determined from intact MS when the probe was added at different molar excesses.** Data are shown as mean and standard deviation of three replicate measurements.

| | 10x | | 20x | | 30x | | 40x | | 50x | |
|---|---|---|---|---|---|---|---|---|---|---|
| **Modifications** | **Mean** | **SD** | **Mean** | **SD** | **Mean** | **SD** | **Mean** | **SD** | **Mean** | **SD** |
| **0** | 3.987 | 3.460 | 0 | 0 | 0 | 0 | 0 | 0 | 0 | 0 |
| **1** | 15.983 | 7.152 | 3.306 | 5.726 | 1.741 | 3.015 | 0 | 0 | 0 | 0 |
| **2** | 24.569 | 5.511 | 12.383 | 7.016 | 7.215 | 9.735 | 2.695 | 4.668 | 0 | 0 |
| **3** | 24.087 | 0.479 | 19.162 | 4.969 | 13.117 | 13.767 | 5.420 | 5.817 | 0 | 0 |
| **4+** | 31.374 | 15.702 | 65.149 | 17.605 | 77.927 | 26.472 | 91.885 | 10.479 | 100 | 0 |

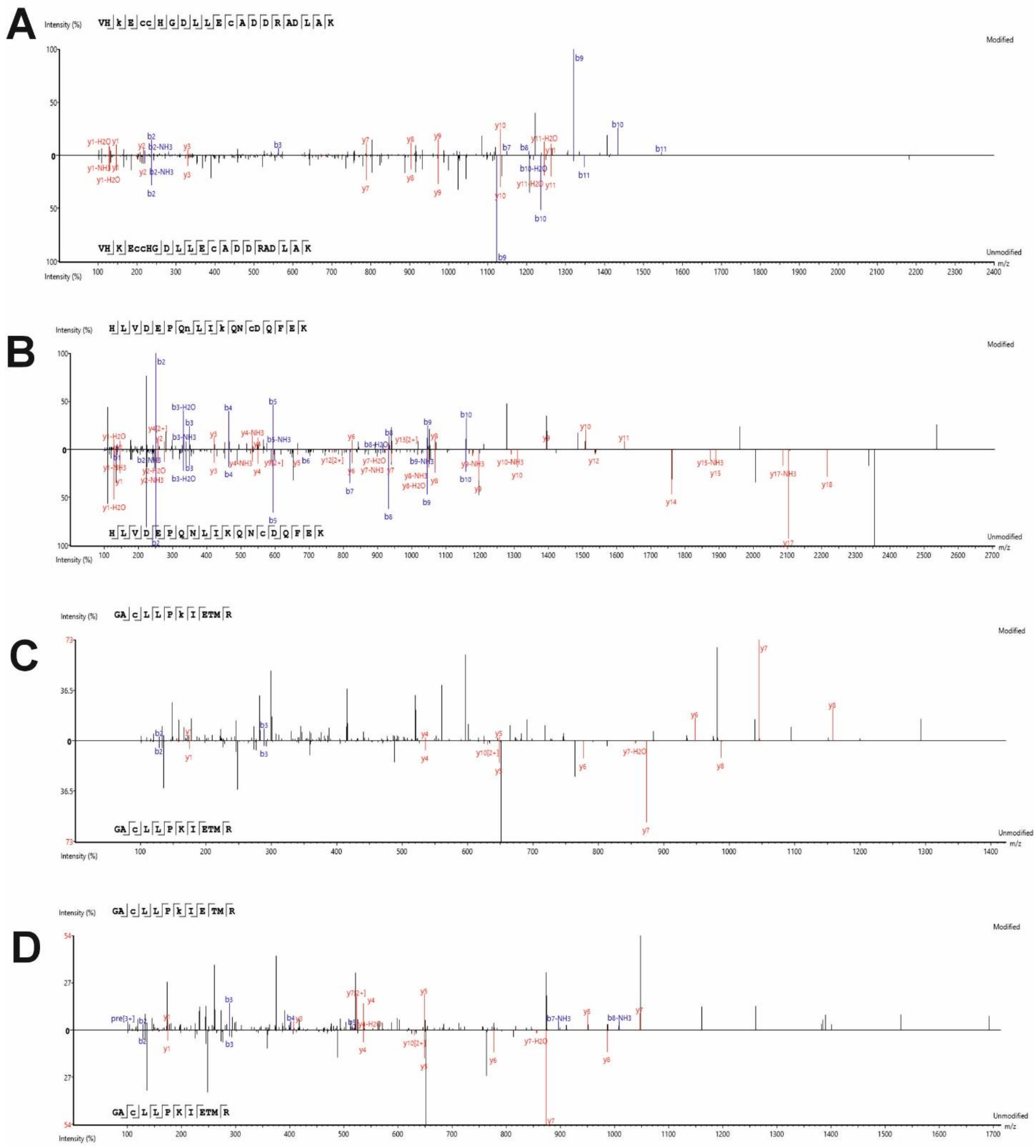


**Supplementary Figure 1. Exemplar MS/MS spectra of modified and unmodified peptides from labelling BSA with probe *1a* and *1b*. (A)** MS/MS spectrum of a tryptic peptide from BSA modified with the indazolone product using probe ***1a***. The MS/MS spectrum of the corresponding unmodified peptide is shown below. **(B)** MS/MS spectrum of a tryptic peptide from BSA modified with the secondary amine product using probe ***1a***. The MS/MS spectrum of the corresponding unmodified peptide is shown below. **(C)** MS/MS spectrum of a tryptic peptide from BSA modified with the indazolone product using probe ***1b***. The MS/MS spectrum of the corresponding unmodified peptide is shown below. **(D)** MS/MS spectrum of a tryptic peptide from BSA modified with the secondary amine product using probe ***1b***. The MS/MS spectrum of the corresponding unmodified peptide is shown below.

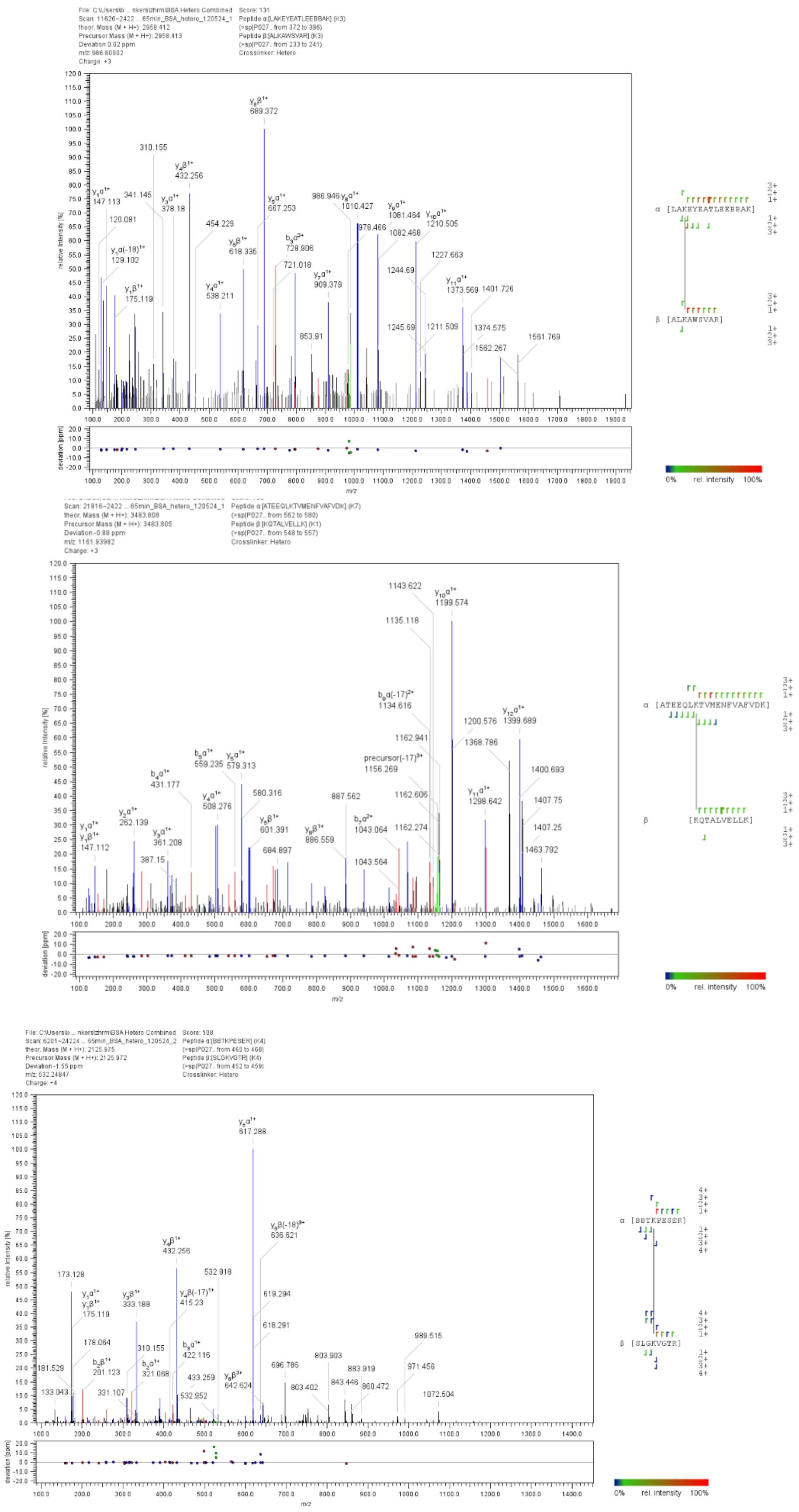


**Supplementary Figure 2. Exemplar MS/MS spectra of identified crosslinks using crosslinker *3* with BSA.**

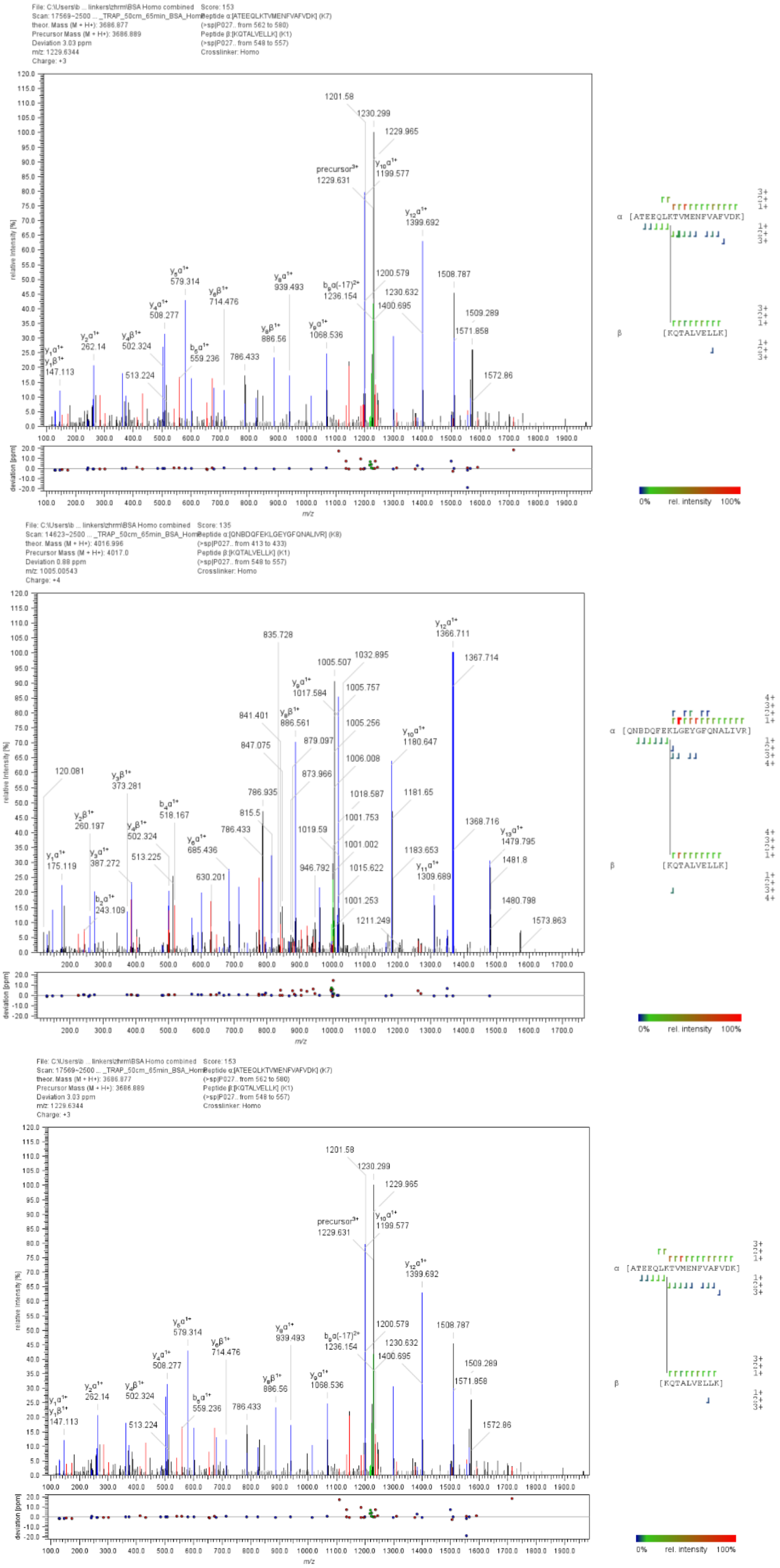


**Supplementary Figure 3. Exemplar MS/MS spectra of identified crosslinks using crosslinker *4* with BSA.**

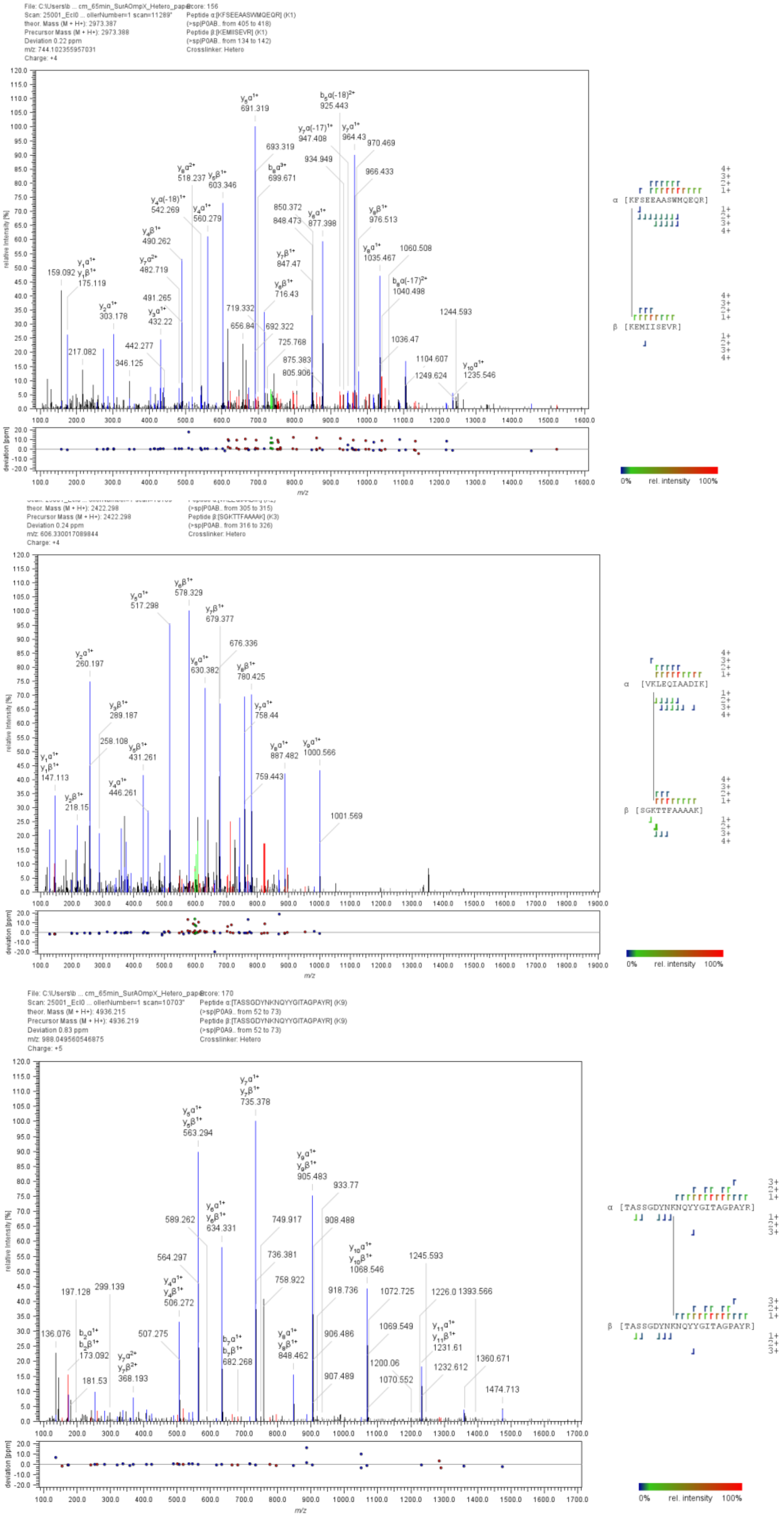


**Supplementary Figure 4. Exemplar spectra of identified crosslinks using crosslinker *3* with SurA-OmpX.**

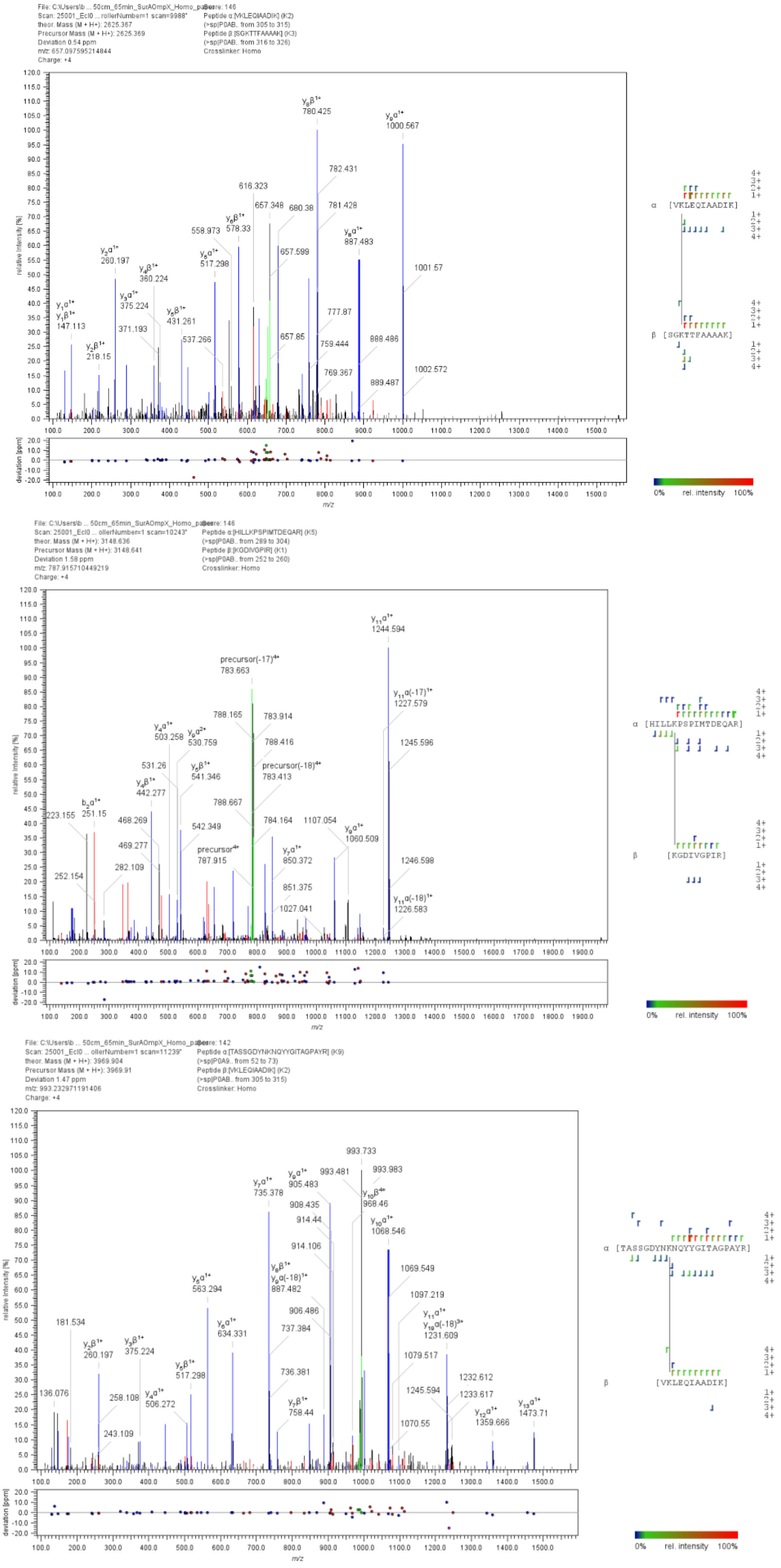


**Supplementary Figure 5. Exemplar spectra of identified crosslinks using crosslinker *4* with SurA-OmpX.**

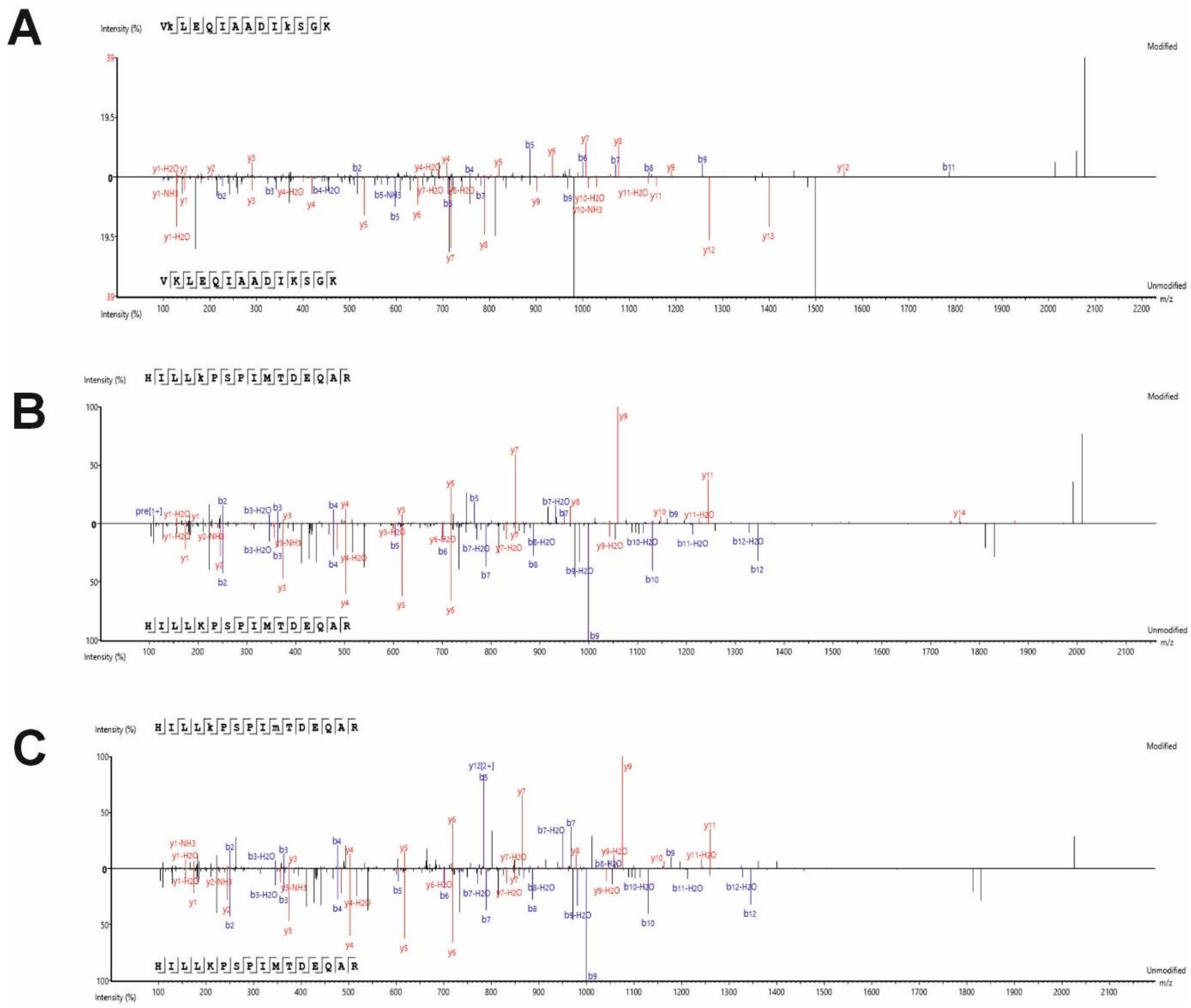


**Supplementary Figure 6. Exemplar MS/MS spectra of mono-linked species and unmodified peptides from crosslinking SurA-OmpX using crosslinker *3*. (A)** MS/MS spectrum of a tryptic peptide modified with the lysine quench product using crosslinker ***3***. The MS/MS spectrum of the corresponding unmodified peptide is shown below. **(B)** MS/MS spectrum of a tryptic peptide modified with crosslinker ***3*** (and no quenching). The MS/MS spectrum of the corresponding unmodified peptide is shown below. **(C)** MS/MS spectrum of a tryptic peptide modified with the hydrolysed quench product using crosslinker ***3***. The MS/MS spectrum of the corresponding unmodified peptide is shown below.

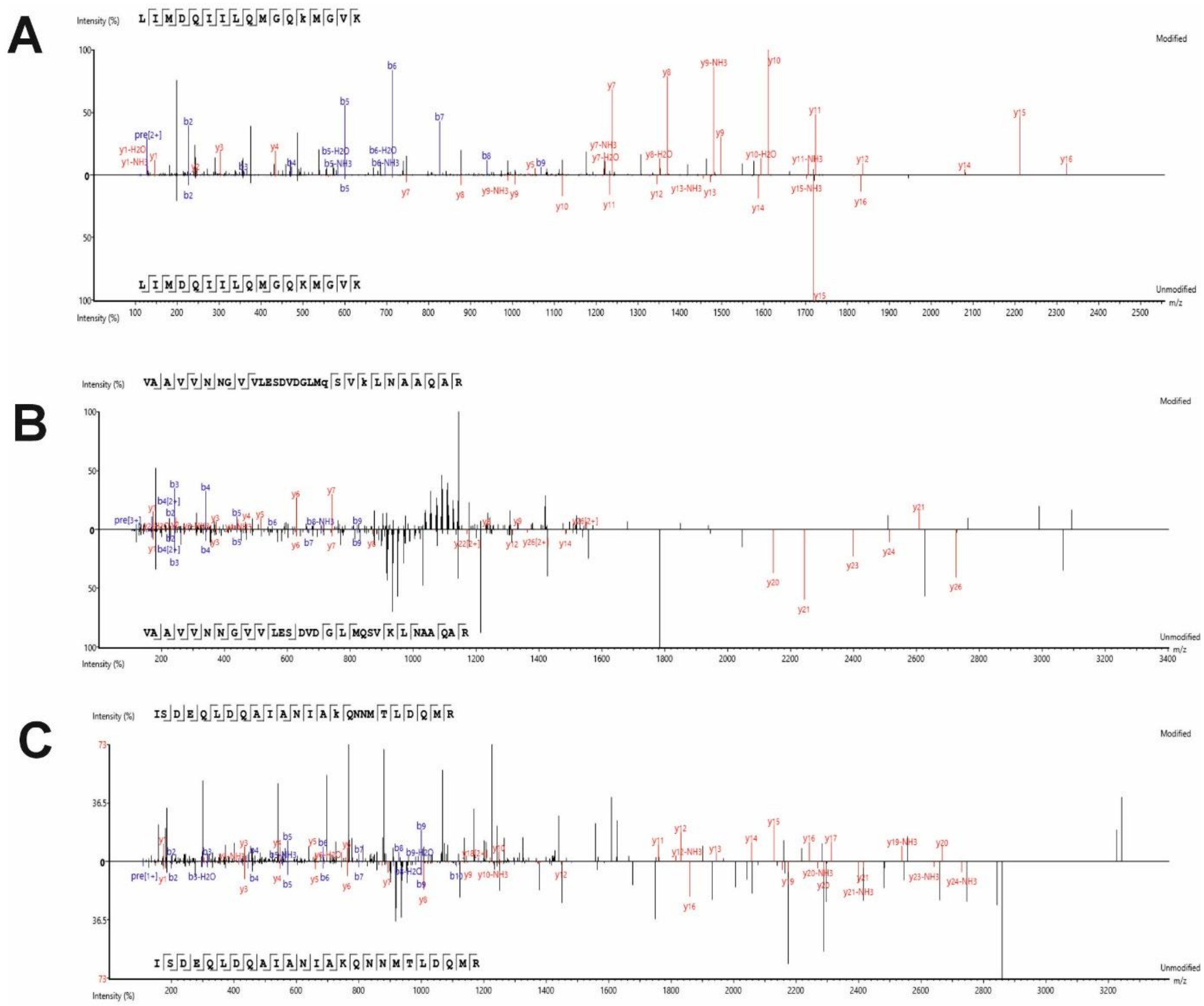


**Supplementary Figure 7. Exemplar MS/MS spectra of mono-link species and unmodified peptide from crosslinking SurA-OmpX using crosslinker *4*. (A)** MS/MS spectrum of a tryptic peptide modified with the lysine quench product using crosslinker ***4***. The MS/MS spectrum of the corresponding unmodified peptide is shown below. **(B)** MS/MS spectrum of a tryptic peptide modified with crosslinker ***4*** (and no quenching). The MS/MS spectrum of the corresponding unmodified peptide is shown below. **(C)** MS/MS spectrum of a tryptic peptide modified with the hydrolysed quench product using crosslinker ***4***. The MS/MS spectrum of the corresponding unmodified peptide is shown below.
